# Two classes of active transcription sites and their roles in developmental regulation

**DOI:** 10.1101/2020.06.23.167338

**Authors:** Sarah Robinson-Thiewes, John McCloskey, Judith Kimble

**Affiliations:** University of Wisconsin-Madison, Department of Genetics; University of Wisconsin-Madison, Department of Biochemistry

## Abstract

Genes encoding powerful developmental regulators are exquisitely controlled, often at multiple levels. Here, we use single molecule FISH (smFISH) to investigate nuclear active transcription sites (ATS) and cytoplasmic mRNAs of three key regulatory genes along the *C. ele*gans germline developmental axis. The genes encode ERK/MAP kinase and core components of the Notch-dependent transcription complex. Using differentially-labeled probes spanning either a long first intron or downstream exons, we identify two ATS classes that differ in transcriptional progression: iATS harbor partial nascent transcripts while cATS harbor full-length nascent transcripts. Remarkably, the frequencies of iATS and cATS are patterned along the germline axis in a gene-, stage- and sex-specific manner. Moreover, regions with more frequent iATS make fewer full-length nascent transcripts and mRNAs, whereas those with more frequent cATS produce more of them. We propose that the regulated balance of these two ATS classes has a major impact on transcriptional output during development.

## Introduction

The control of gene expression is central to animal development and homeostasis. To achieve that control, an increasingly complex choreography of regulatory steps dictates when, where, and how much mRNA is produced. Transcriptional initiation has taken center stage as the key step in regulating gene expression for years (Mannervik et al., 1999), but other downstream mechanisms have now joined initiation on that stage. Most relevant to this work is regulation of transcriptional progression. A classic example of regulation at this step occurs in *Drosophila* embryos, where the cell cycle is too short to complete transcription through unusually long Hox genes before the cell divides (Gubb, 1986; Shermoen and O’Farrell, 1991). A more broadly used mechanism is the regulated release from transcriptional pauses that occur ∼20-60 bp after initiation, the “promoter-proximal pause” (Adelman and Lis, 2012). Transcriptional progression can be deduced with “-omic” methods, such as PRO-seq or NET-seq (Churchman and Weissman, 2011; Jonkers and Lis, 2015), or with imaging methods, such as single molecule fluorescence *in situ* hybridization (smFISH) or live-imaging (Pichon et al., 2018). One advantage of smFISH is that transcription of endogenous genes can be followed in their native context with single cell and single molecule resolution. Although this work began with an smFISH investigation of active transcription sites (ATS) and transcriptional yields at key regulatory genes during development, it led to discovery of two distinct ATS classes and what we propose is a case of regulated transcriptional progression.

Our studies have been conducted in the adult *C. elegans* gonad, which is well-suited to smFISH (e.g. Lee et al., 2016) and where germ cell development occurs linearly along the distal-proximal axis. Germline stem cells (GSCs) reside within their niche at the distal end and GSC daughters mature progressively toward gametogenesis as they move from the niche and ultimately reach the other end (Figure 1A). Previous studies in this tissue revealed a gradient of Notch-dependent transcriptional bursts at the Notch target gene, *sygl-1* (Crittenden et al., 2019; Lee et al., 2019, 2016). Here, we focus on transcription of three other genes that regulate GSC self-renewal or differentiation (Figure 1B). The *lag-1* and *lag-3* genes encode core components of the Notch transcriptional activation complex, which promotes GSC self-renewal in response to signaling from the niche (Christensen et al., 1996; Lambie and Kimble, 1991; Petcherski and Kimble, 2000). The *mpk-1* gene encodes ERK/MAP kinase (Lackner et al., 1994; Wu and Han, 1994), which promotes several aspects of germline differentiation: sperm/oocyte fate specification (Min Ho Lee et al., 2007; Morgan et al., 2013), progression through meiotic prophase, and oogenesis (Arur et al., 2011; Church et al., 1995; Lopez 3rd et al., 2013). We emphasize that our approach queries transcription at endogenous genes in wildtype animals; our results therefore avoid potentially confounding effects of transgenes, inserted tags, and reporter constructs.

**Figure 1:**
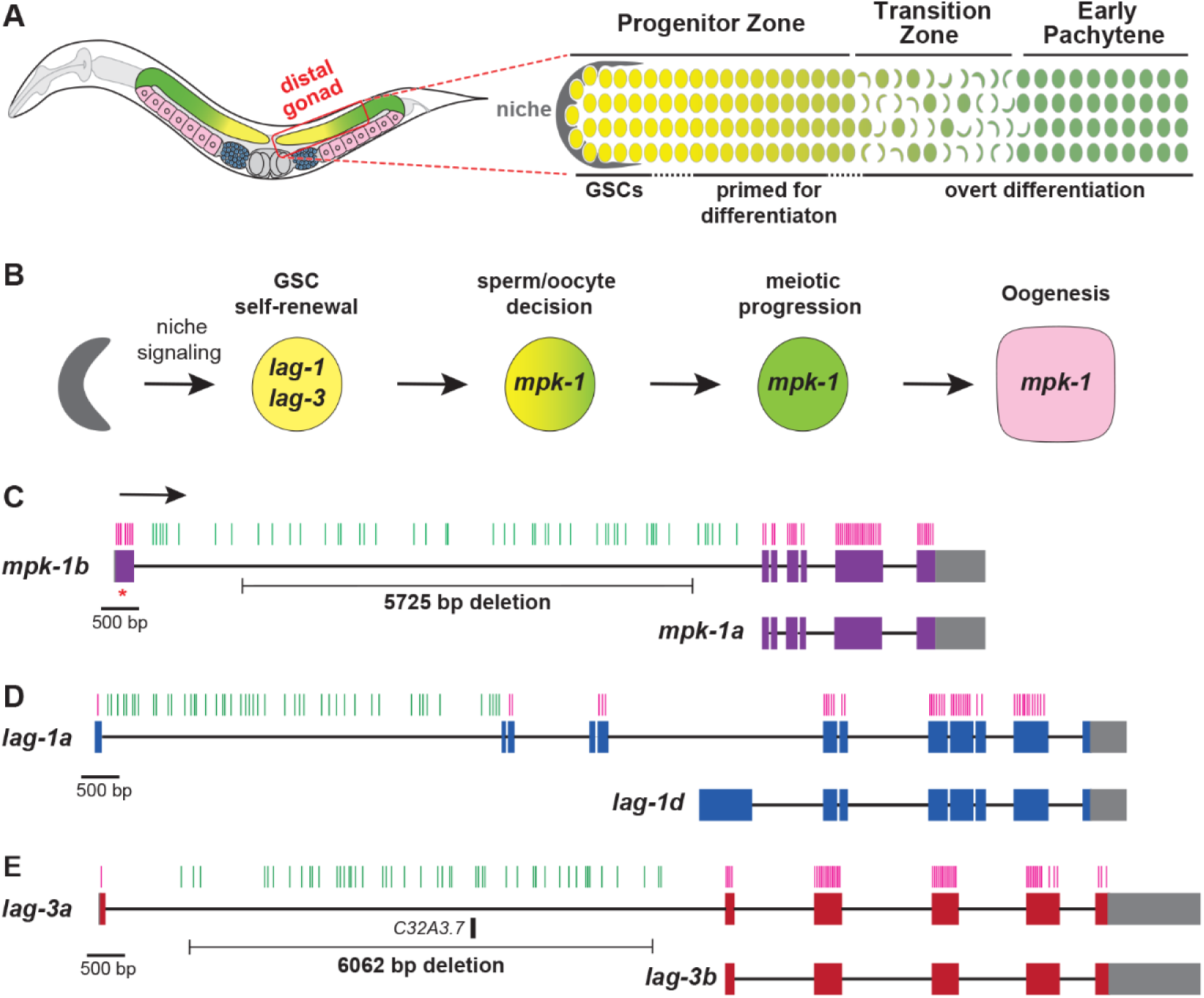
*C. elegans* germline anatomy and key regulatory genes. **A**. Left, adult hermaphrodite with two U-shaped gonadal arms (in color) that consist largely of germ cells maturing along a distal-proximal axis. In the distal gonad (red square), germline progenitors divide mitotically (yellow) and then enter meiotic prophase (green); in the proximal gonad, germ cells differentiate into sperm (blue) or oocytes (pink). Right, distal gonad. A single-celled somatic niche (grey) maintains germ cells in a naïve stem cell state (GSCs). The Progenitor Zone (PZ) includes GSCs within the niche and their daughters primed to differentiate; more proximally, germ cells enter meiotic prophase (green crescents). Transitions between germ cell states are marked as dashed lines. **B**. Regulators relevant to this work. Color scheme as in A. The *lag-1* and *lag-3* genes encode niche signaling components, the LAG-1 CSL DNA binding protein and LAG-3 Mastermind-like transcription factor (Christensen et al., 1996; Petcherski and Kimble, 2000); *mpk-1* encodes MPK-1 ERK/MAP kinase, which promotes sperm fate specification, meiotic progression, and oogenesis (Arur et al., 2011; Church et al., 1995; Min Ho Lee et al., 2007; Lopez 3rd et al., 2013; Morgan et al., 2013). **C-E**. Architecture of genes encoding key regulators. Boxes, exons; lines, introns. Gene-specific colors in exons denote coding regions and grey indicates untranslated regions (UTR). The direction of transcription is the same for all genes (arrow in C). Short vertical lines above genes indicate sites of individual probes in the probe sets used for smFISH; these probe sets target either the large first intron (green) or all exons (magenta). Red asterisk marks site of a 1 bp frame-shifting insertion in the *mpk-1b* first exon, a mutant used as a control for *mpk-1* exon probe specificity. Deletions in *mpk-1* and *lag-3* long first introns were used to test for intron probe specificity. The *lag-1* gene makes four isoforms (*lag-1a-d*); *lag-1a-c* differ in size of exons 2 and 3 and for simplicity, *lag-1a* is shown to represent *lag-1a-c*. The *lag-3a* first intron contains a predicted ncRNA *C32A3*.*7*.

To obtain a high resolution and quantitative view of transcription during development, we coupled smFISH with a MATLAB image analysis code to score RNAs with 3D resolution in the germline tissue, as done previously for *sygl-1* (Crittenden et al., 2019; Lee et al., 2016). Using differentially labeled probe sets to either the 5’ half to two-thirds of each gene (spanning the long first intron) or the remaining 3’ part (the exons, most of which are 3’ to the intron), we found two distinct ATS classes that differ in transcriptional progression. One class, iATS for “incomplete” ATS, harbors partial nascent transcripts; the other class, cATS for “complete” ATS, harbors full length nascent transcripts. Remarkably, the frequencies of these ATS classes are gene-, position- and sex-specific, suggesting developmental regulation. Most strikingly, iATS and cATS frequencies are reciprocally graded for two genes in the stem cell region, in a manner consistent with the graded expression of those genes. We propose that regulated changes in transcriptional progression, inferred from changes in the frequency of ATS class, drives the developmental expression of these genes.

## RESULTS

### Experimental design and a modified MATLAB code

Our experimental design took advantage of the simple architecture of the *C. elegans* adult germline (Figure 1A), the ability to visualize transcription at high resolution in this tissue using smFISH (Crittenden et al., 2019; Lee et al., 2016), and genes with long first introns that encode key regulators of germline development (Figure 1B). The *mpk-1, lag-1*, and *lag-3* first introns are 8.2 kb, 6 kb, and 8.5 kb, respectively. We used Stellaris® Probe Designer to create two probe sets for each gene, spanning either the long first intron and thus covering the 5’ half to two-thirds of each gene or all exons and thus covering the remaining 3’ part of the gene (Figure 1C-E). Each set included 47-48 individual probes (see Table S1 for details). The intron and exon probe sets were labeled with different fluorophores to generate distinct signals (Figure 2A-C; Figure S1A; S2A; S3A). Probe specificities were confirmed by inhibiting transcription with α-amanitin to ensure detection of RNA rather than DNA (Figure S1B; S2B; S3B); intron-specific deletions to ensure specificity of the intron probe signal (*mpk-1* and *lag-3* only, an analogous *lag-1* deletion could not be isolated) (Figure S1C, S3C); and a frame-shift mutant (*mpk-1*) or RNAi (*lag-1, lag-3*) to ensure specificity of the exon probe signal (Figure S1D, S2C-D, S3D-E). See Figure S1-3 legends for gene-specific details of these specificity assays.

**Figure 2:**
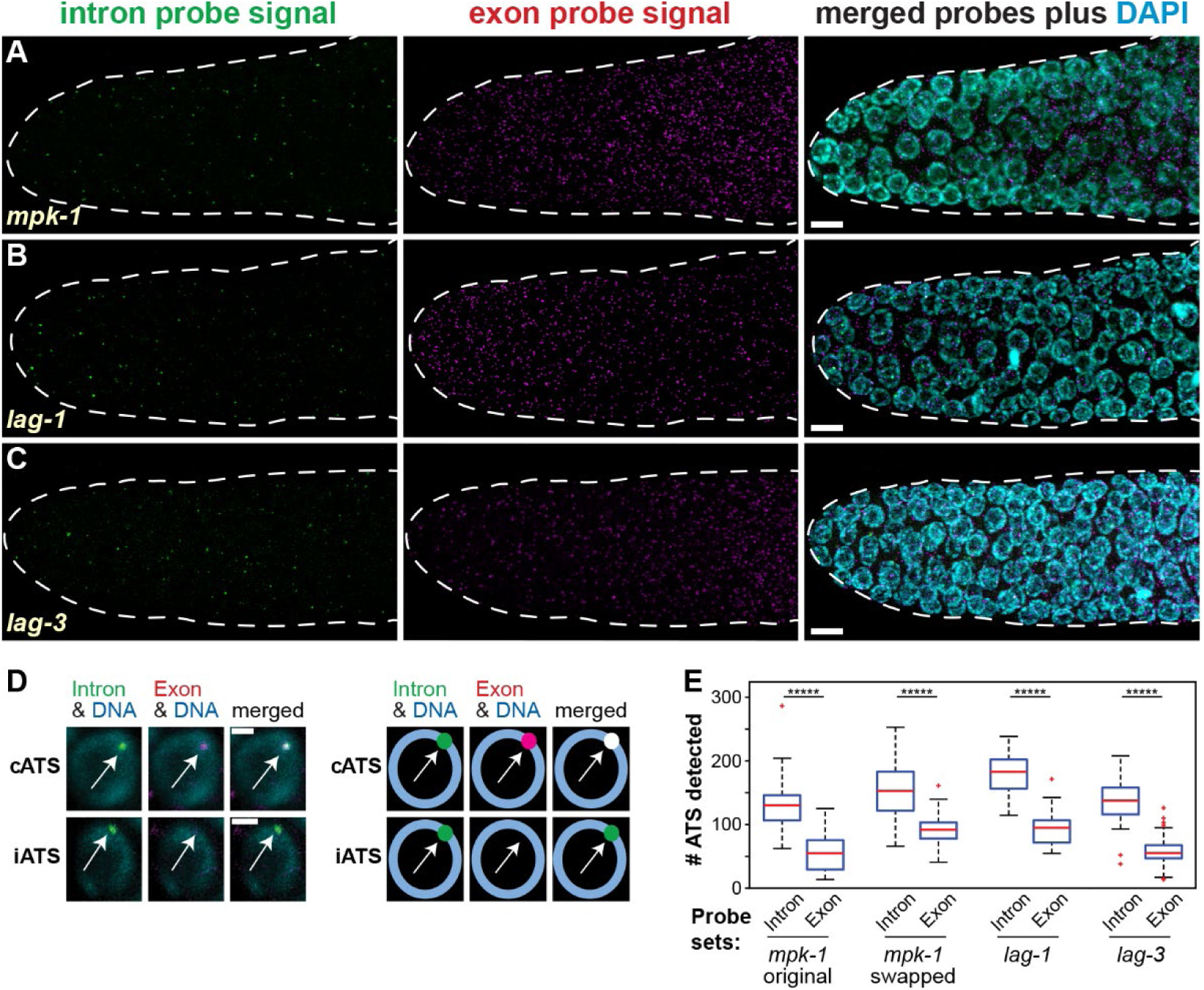
Identification of two types of active transcription sites. **A-C**. Maximum projections of smFISH images. Left, intron probe set signals (green); middle, exon probe set signals (magenta); merge of both signals with DAPI (cyan). Scale bar = 5 µm. **D**. Two classes of active transcription sites (ATS). Left, images; right, cartoons. A cATS (complete ATS) is seen by overlapping intron and exon signals; an iATS (incomplete ATS) is seen by a unique intron signal. **E**. ATS numbers, regardless of type, detected with either the intron probe set (intron) or the exon probe set (exon) for each gene. Overlaps of the intron-detected and exon-detected spots were not determined in this analysis. To test if high intron detection reflected fluorophore bias, the fluorophores conjugated to the original *mpk-1* intron and exon probes were swapped. *****p<0.0000001, Student’s t-test.

A previously published MATLAB code (used to analyze *sygl-1* transcription in the *C. elegans* progenitor zone) defined ATS as nuclear spots with overlapping exon and intron probe signals (Lee et al., 2016). However, a preliminary inspection by eye of the smFISH images for *mpk-1, lag-1*, and *lag-3* transcripts revealed two types of nuclear spots, both with morphology and size of an ATS. We categorize these two types as “cATS” and “iATS” (Figure 2D). cATS are detected with overlapping signals from the exon and intron probe sets, while iATS are detected uniquely with the intron probe set. The finding of these two ATS types caused us to modify the original code. Briefly, the new code detects nuclear signals independently for the exon and intron probe sets, it determines if the two signals are overlapping to assign them as cATS or iATS, and it identifies cytoplasmic mRNA, all within the 3D gonad (Figure S4, see Methods). During this code revision, we asked how many nuclear spots were seen with each probe set, without taking into consideration any overlap, and found that the intron probe detected more spots than the exon probe for all three genes (Figure 2E). To test whether the abundance bias of intron spots might reflect fluorophore differences, we tested *mpk-1* probe sets with swapped fluorophores, but the intron spot abundance bias did not change (Figure 2E). To validate our modified MATLAB code, we tested it by rescoring previously published images of *sygl-1* smFISH and obtained results equivalent to previous report (Figure S5) (Lee et al., 2016). The new code thus scores cATS and iATS with spatial resolution.

### Transcriptional probabilities and yields in the progenitor zone

We analyzed *mpk-1, lag-1*, and *lag-3* transcription in the adult hermaphrodite progenitor zone (PZ), where GSCs reside distally and more proximal GSC daughters become primed for differentiation (Cinquin et al., 2010; Crittenden et al., 2006; Rosu and Cohen-Fix, 2017). Position within the PZ was scored as the number of germ cell diameters (gcd) or “rows” along the distal-proximal axis from the distal end, by convention. For this study, we focused on the distal-most 12 rows of the PZ, a region with roughly 150 germ cells that divide asynchronously every ∼6 hrs on average (Albert Hubbard and Schedl, 2019) and move proximally at a rate of ∼0.4 to 1 row per hour (Rosu and Cohen-Fix, 2017). Importantly, a germ cell’s state — naïve and stem cell-like or triggered to differentiate — corresponds to its position along the axis (Figure 3A).

**Figure 3:**
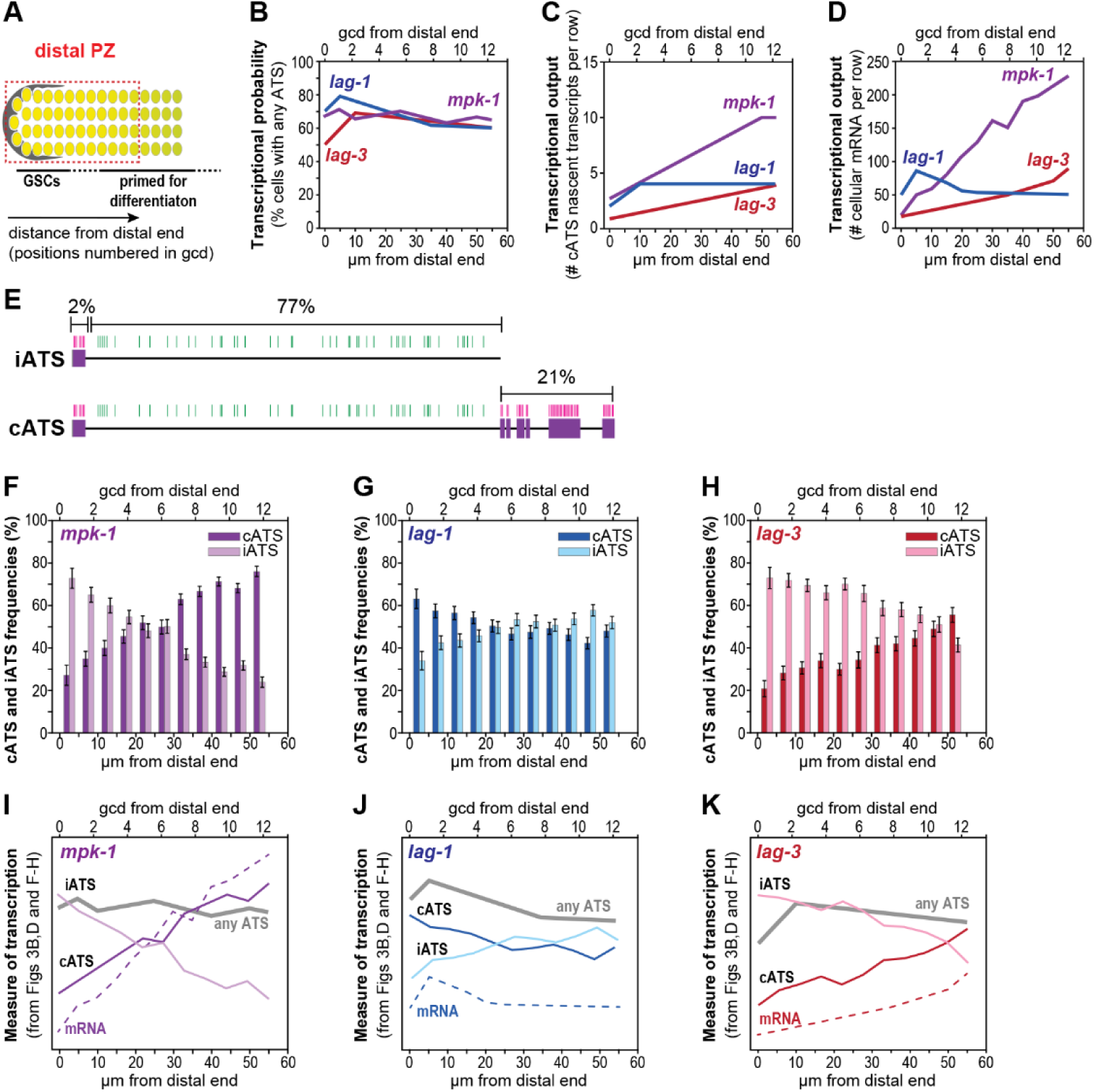
Transcription of *mpk-1, lag-1* and *lag-3* in the distal progenitor zone. **A**. Gonadal region scored (dashed red box) extends 12 germ cell diameters (gcd) into the progenitor zone (PZ) from the niche (grey). Numbers indicate position as gcd along the distal-proximal axis from the distal end, according to convention; MATLAB scores position in µm from the distal end, with each gcd averaging ∼4.4 µm in this region (Lee et al., 2016). **B-D and F-K**. x-axis represents position in gcd (top) and μm (bottom). Number of gonads scored for each gene: *mpk-1*, n = 37; *lag-1*, n = 36; and *lag-3*, n = 32. **B**. Transcriptional probability measured as percentage of cells with at least one ATS, including iATS and cATS. Line plots derived from data in Figure S6(A-C). Total number of cells scored: *mpk-1*, n = 6065; *lag-1*, n = 5981; and *lag-3*, n = 4472. **C**. Transcriptional output measured as total number of nascent transcripts at cATS, per cell row. We limited analysis to nascent transcripts at cATS, as explained in text and detailed in Methods. Line plots derived from data in Figure S6(G-I). **D**. Transcriptional output measured as total number of cellular mRNAs per cell row; mRNAs in rachis were excluded. Germ cell boundaries were determined from MATLAB-generated Voronoi cells as described in Methods. Line plots derived from data in Figure S6(J-L). Data for number of mRNA per cell in Figure S6(M-O). **E**. Detection of RNAs at iATS (above) and cATS (below). iATS are seen uniquely with the intron probe set whereas cATS are seen with overlapping exon and intron probe sets. The intron probe set spans 77% of the full-length transcript (excluding 3’UTR); by contrast, the exon probe set spans only 23%. **F-H**. iATS and cATS frequencies as a function of position. Each bar shows percentage of total ATS that are cATS (darker bars) or iATS (lighter bars). Numbers of total ATS (cATS plus iATS) scored: *mpk-1*, n = 5699; *lag-1*, n = 6610; and *lag-3*, n = 4200. Standard errors are shown. **I-K**. Measures of transcription taken from panels above and combined to highlight patterns of graded increase, decrease, or relative uniformity; individual lines represent quite different measures and specific values are therefore not comparable. Transcriptional probability (gray line) from panel 3B; cellular mRNA abundance per row (dashed line) from panel 3D; cATS frequency (dark colored line) and iATS frequency (lighter colored line) from panels 3G-I. See original panels for y-axis data ranges.

We first determined the percentage of germ cells harboring any ATS, cell row by cell row along the PZ developmental axis. The ATS scored in this initial analysis include both cATS and iATS, and thus reveal genes that have not only initiated transcription but also elongated far enough for detection with smFISH probes. The percentage of cells with any ATS therefore provides a measure of transcriptional probability, as established previously (Chubb et al., 2006; Lee et al., 2019, 2016). The *mpk-1, lag-1*, and *lag-3* genes were all transcribed actively across the distal PZ with 60 to 70% of cells possessing either a cATS or iATS (Figure 3B; Figure S6A-C). For comparison, the Notch-activated *sygl-1* gene produces ATS in ∼65% of the distal germ cells within the niche but <5% outside the niche, as previously reported (Lee et al., 2016) and confirmed here (Figure S5B). We also scored how many individual ATS were seen in each nucleus as a function of position along the axis. As expected for genes that transcribe in bursts within an actively dividing cell population, the numbers of individual ATS per cell varied between zero and four (Figure S6D-F). The higher percentage in the most distal germ cells for *lag-1* ATS and lower percentage in the same region for *lag-3* ATS are reproducible but not understood. Most importantly, all three genes are actively engaged in transcription across the distal PZ.

We next scored two measures of transcriptional productivity as a function of position in the PZ—the number of nascent transcripts at cATS (Figure 3C, Figure S6G-I) and number of mRNAs in the cytoplasm (Figure 3D, S6J-O). Measurement of nascent transcript abundance was limited to cATS, because it could be estimated by comparing intensity values of exon probe spots at cATS in the nucleus and exon probe dots at single mRNAs in the cytoplasm. While this strategy misses nascent transcripts at iATS, the compared values rely on the same probe, same fluorophore, and same image. Both the number of *mpk-1* and *lag-3* nascent transcripts increased steadily as germ cells moved along the PZ axis, while the number of *lag-1* nascent transcripts increased initially and then leveled off (Figure 3C). Following those trends, *mpk-1* and *lag-*3 mRNA numbers also increased along the axis, while numbers of *lag-1* mRNAs were more level (Figure 3D). The pattern of *lag-1* mRNA abundance conforms for the most part with an independent report published recently (Chen et al., 2020). Most strikingly, the nearly uniform transcriptional probabilities for *mpk-1* and *lag-3* (Fig 3B) did not match their gradually increasing transcriptional outputs across the PZ axis (Figs 3C, 3D).

### Frequency of one ATS class, the cATS, corresponds to mRNA yield

Why might transcriptional probability and output have distinct patterns for a given gene? We considered the possibility that the frequency of the two ATS classes, iATS and cATS (Figure 2), underlie the explanation. iATS are uniquely detected with the first intron probe, so nascent transcripts at iATS have elongated through much of the long intron, but not the downstream exons (Figure 3E, top). By contrast, cATS are detected with both exon and intron probes, so nascent transcripts at cATS must have elongated through both the long intron and the downstream exons (Figure 3E, bottom). Because iATS and cATS reveal transcription sites dominated by different extents of transcriptional progression, cATS would be expected to yield a more robust transcriptional output than iATS. Based on that idea, we determined the frequencies of each ATS class, measured as a percentage of all ATS, and asked if their frequencies change along the PZ developmental axis. One might have thought that iATS and cATS frequencies would simply reflect the extent of sequence covered by each probe set. If that were the case, they would be the same regardless of position. However, iATS and cATS frequencies had gene-specific patterns along the PZ axis (Figure 3F-H). For example, *mpk-1* cATS frequency increased steadily from ∼30% distally to ∼75% at row 12, and iATS frequency decreased correspondingly (Figure 3F). A similar trend was found for *lag-3* (Figure 3H), but the *lag-1* cATS frequency initially decreased and then leveled off across the PZ (Figure 3G). These patterns suggest that transcriptional progression is gene-specific and changes as germ cells move through the PZ. A logical extension of that idea is that the pattern of only one class, the cATS with their full or nearly full-length transcripts, is responsible for the pattern of mRNA production. That prediction was borne out by comparing patterns of ATS, cATS, iATS, and mRNAs in gene-specific graphs (Figure 3I-J) and finding positive correlations between cATS frequency and mRNA number and negative correlations between iATS frequency and mRNA number (Table 1). We conclude that cATS frequency, not iATS frequency, drives mRNA output as germ cells mature through the PZ (see Discussion).

**Table 1:** Pearson’s correlation tests between measures of transcription

**A. Transcriptional probability (% cells with any ATS) vs transcriptional output (# mRNA in cells)**
| Gene | Correlation coefficient (r) | p value |
| --- | --- | --- |
| *mpk-1* | 0.0157 | 0.7456 |
| *lag-1* | 0.0725 | 0.1362 |
| *lag-3* | -0.0594 | 0.0594 |

**B. cATS frequency (% ATS that are cATS) vs transcriptional output (# mRNA in cells)**
| Gene | Correlation coefficient (r) | p value |
| --- | --- | --- |
| *mpk-1* | 0.4872 | $7.5725 \times 10^{-28}$ |
| *lag-1* | 0.1272 | 0.0081 |
| *lag-3* | 0.4448 | $4.6640 \times 10^{-20}$ |

**C. iATS frequency (% ATS that are iATS) vs transcriptional output (# mRNA in cells)**
| Gene | Correlation coefficient (r) | p value |
| --- | --- | --- |
| *mpk-1* | -0.4520 | $9.6179 \times 10^{-24}$ |
| *lag-1* | -0.1149 | 0.0169 |
| *lag-3* | -0.3836 | $6.5252 \times 10^{-15}$ |

**D) Transcriptional output (# nascent transcripts) vs transcriptional output (# mRNA in cells)**
| Gene | Correlation coefficient (r) | p value |
| --- | --- | --- |
| *mpk-1* | 0.8214 | $1.9340 \times 10^{-106}$ |
| *lag-1* | 0.3807 | $2.3929 \times 10^{-16}$ |
| *lag-3* | 0.8074 | $3.9855 \times 10^{-86}$ |

### mpk-1, lag-1, and lag-3 transcription in the early Pachytene Region

We next investigated transcription of the same three genes in a different region of the gonad, where germ cells have begun to differentiate. Specifically, we focused on 12 rows of germ cells that begin at the proximal boundary of the Transition Zone (TZ) and extend into the Pachytene Region (Figure 4A). Germ cells here have entered the pachytene stage of meiotic prophase and begun oogenesis. For simplicity, we refer to the region as EP for early pachytene. Though germ cells are not dividing mitotically, they move proximally through the EP at a rate of ∼1 row per hour (Albert Hubbard and Schedl, 2019; Crittenden et al., 2006), and progressively mature as they move along the distal-proximal axis.

**Figure 4:**
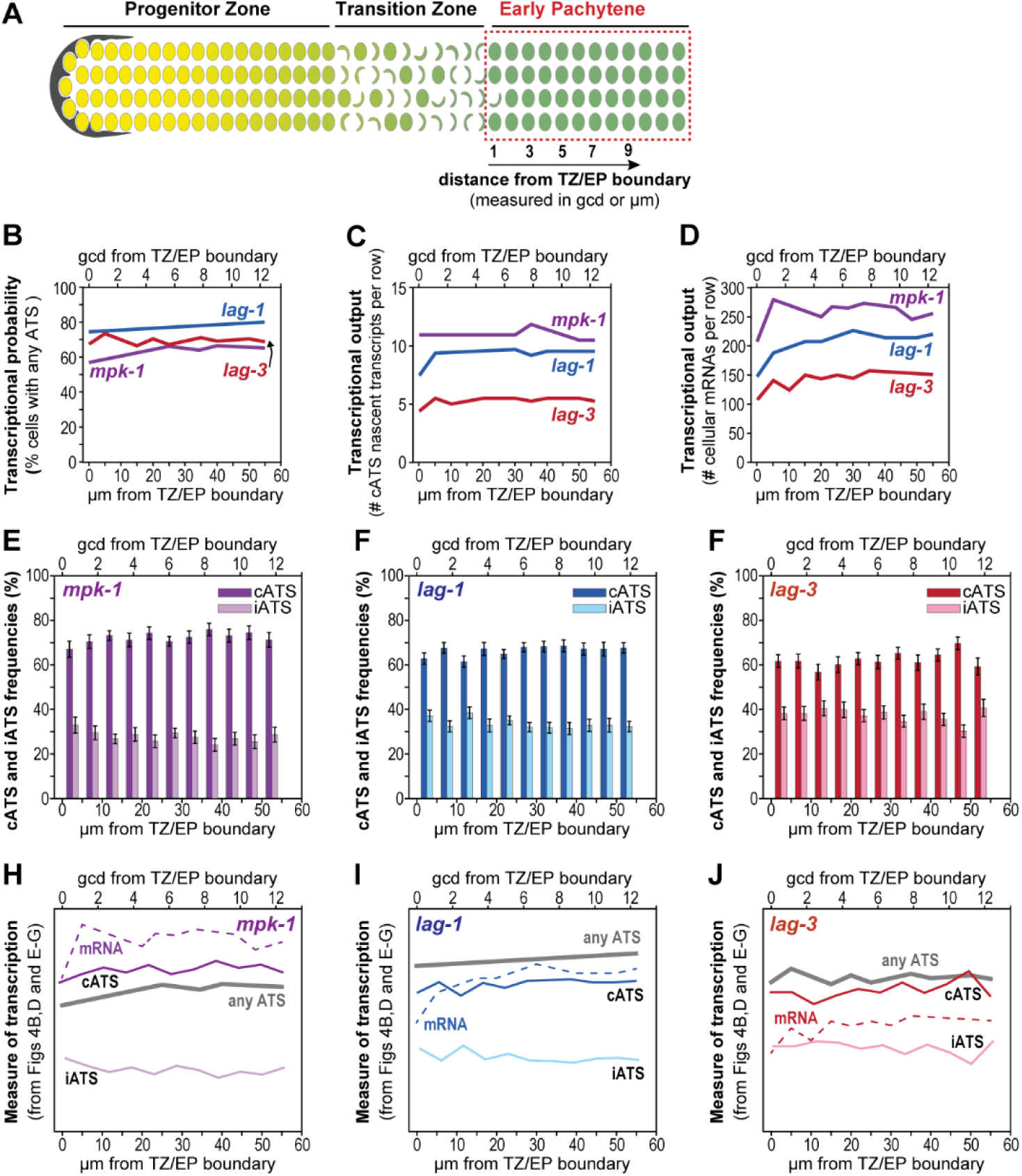
Transcription of *mpk-1, lag-1* and *lag-3* in the Early Pachytene region. **A**. Gonadal region scored in this figure (dashed red box) extends 12 gcd into the early pachytene region (EP) from the TZ/EP boundary. Numbers mark position or cell row, measured in gcd and starting at the TZ/EP boundary; MATLAB scores position in µm from the same boundary with each gcd averaging ∼4.4 µm in this region. See Figure S7 for smFISH images. **B-J**. Quantification of transcripts as a function of position, per cell row. Number of gonads analyzed for each gene: *mpk*-1, n = 36; *lag-1*, n = 36; and *lag-3*, n = 37. x-axis represents cell position as gcd (top) and μm (bottom) from the TZ/EP boundary. **B**. Percent cells with one or more ATS, either cATS or iATS. Total cell number scored: *mpk-1*, n = 6249; *lag-1*, n =6064; and *lag-3*, n = 4625. Line plots from data in Figure S8(A-C). **C**. Transcriptional output measured as total nascent transcripts at cATS per cell row (see text and Methods). Line plots from data in Figure S8(G-I). **D**. Transcriptional output measured as total number of mRNAs in cells per row; mRNAs in rachis were excluded. Cells were defined as described in Figure 2D. Line plots from data in Figure S8(J-L). Data for number of mRNA per cell in Figure S8(M-O). **E-G**. iATS and cATS frequencies as a function of position. Bars show percentages of total ATS that are cATS (darker bars) or iATS (lighter bars). Total number of ATS (either cATS or iATS) scored: *mpk-1*, n = 5749; *lag-1*, n = 10118; and *lag-3*, n = 4730. **H-J**. Measures of transcription taken from panels above and combined to highlight patterns; individual lines represent quite different measures and specific values are therefore not comparable. Transcriptional probability (gray line) from 4B. Number of mRNA in cells (dashed colored line) from 4D. The cATS frequency (dark colored line) and iATS frequency (lighter colored line) from data in 4E-G. See original panels for y-axis data ranges.

Analyses in the EP paralleled those in the distal PZ (see Figure S7 for representative smFISH images). The *mpk-1, lag-1*, and *lag-3* genes were actively transcribed across the region: the percentages of cells with any ATS ranged from ∼50-80% (Figure 4B; Figure S8A-C), and ATS numbered zero to four per nucleus as expected (Figure S8D-F). The number of nascent transcripts at cATS was steady across the region for each gene (Figure 4C; Figure S8G-I), while mRNA numbers were essentially uniform or increased slightly (Figure 4D; Figure S8J-O). Strikingly and in contrast to the PZ, most ATS were cATS throughout the EP region (Figure 4E-G). Display of the various measures of transcription in a single graph highlights their similarity (Figure 4H-J). Again, high cATS frequencies match abundance of cytoplasmic mRNAs. Thus, the ATS class frequencies and hence transcriptional progression appear relatively uniform for these genes as they move through early pachytene.

### Deletions in the *mpk-1* long first intron have either minor or no detectable germline defects

We considered the idea that the size or content of the long first intron of each gene might influence the pattern of cATS frequency in the PZ. Introns can contain regulatory elements that affect numerous aspects of the transcription process, including elongation and splicing (Chorev and Carmel, 2012; Gubb, 1986; Parenteau et al., 2019; Rose, 2019; Swinburne et al., 2008; Swinburne and Silver, 2008; Takashima et al., 2011). To address this issue, we focused on *mpk-1* for two reasons. First, the *mpk-1b* isoform is the principal and likely only *mpk-1* transcript in the germline (Figure 1C) (Lee et al., 2007; Robinson-Thiewes et al., in preparation). As a result, *mpk-1* smFISH in the germline scores this isoform specifically. By contrast, tissue-specificity is unknown for the isoforms of the other genes (Figure 1D-E). Second, a 5.7 kb deletion in the *mpk-1* intron (Figure 1C) is homozygous viable. Therefore, the regulatory effects of potential elements within the *mpk-1* long first intron can be investigated with mutants. By contrast, an analogous deletion in *lag-3* (Figure 1D) is embryonic lethal and the analogous *lag-1* intron deletion could not be recovered.

We made 1 kb deletions at the 5’ end (5’ Δ), middle (mid Δ) and 3’ end (3’ Δ) of the *mpk-1* long first intron at the endogenous locus (Figure 5A). To choose the specific regions removed, we used a combination of homology searches and ChIP datasets. The 5’ Δ mutant removed a 90 bp sequence predicted *in silico* to form a hair-pin loop and is repeated throughout the *C. elegans* genome (Table S2); we chose this region because a hair-pin loop in the first intron of *MYB* attenuates its transcription (Bender et al., 1987; Pereira et al., 2015). The mid Δ mutant removed a region enriched for RNA polymerase (RNAP) II serine 5 phosphorylation and epigenetic markers of an enhancer (Liu et al., 2011); we chose this region because in other genes, intragenic enhancers have been found to attenuate transcription (Cinghu et al., 2017). The 3’ Δ mutant removed a region with no distinguishing features. To avoid splicing defects, the 5’ end of 5’ Δ was placed 287 bp downstream of the 5’ splice site, and the 3’ end of 3’ Δ was placed 170 bp upstream of the predicted branch site and 197 bp upstream of the 3’ splice site.

**Figure 5:**
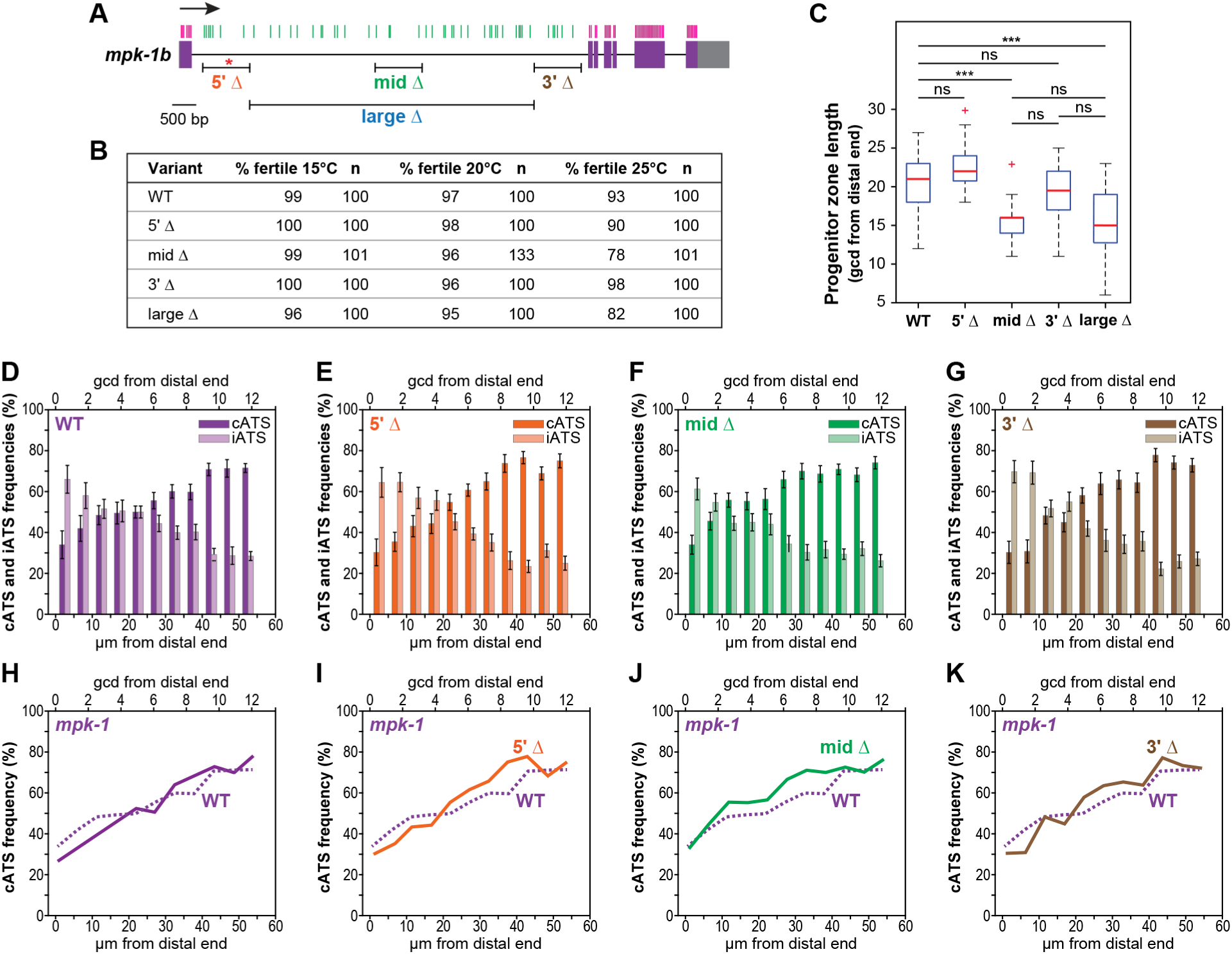
*mpk-1* intron deletion mutants and their effects. **A**. CRISPR-induced deletions in large first intron of the endogenous *mpk-1* locus. Conventions as in Figure 1. Barred lines show extent, position, and name of each deletion. Each smaller deletion is ∼1 kb and the large Δ is ∼ 5.7 kb. Red asterisk marks site of 90 bp motif removed by the 5’ Δ. **B**. Effects of intron deletions on fertility. Mutants were scored for production of embryos at 15°C, 20°C, and 25°C. **C**. Effects of intron deletions on Progenitor Zone length, scored as number of gcd from the distal end to formation of DAPI-stained crescents marking entry into meiotic prophase, by convention. Number PZs counted: WT, n = 22; 5’ Δ, n = 26; mid Δ, n = 12; 3’ Δ, n = 16; large Δ, n = 21. Student’s t-test determined statistical significance. ***, p < 0.00001. **D-K**. iATS and cATS frequencies as a function of position in the Progenitor Zone (dashed red box in Figure 2A). Mutants and wildtype were grown and assayed in parallel under the same conditions; number of individual probe binding sites removed was roughly equivalent for all three small deletions (Table S1). Extended intron data in Figure S9. Number gonads scored: wildtype, n = 16; 5’ Δ, n = 19; mid Δ, n = 20; and 3’ Δ, n = 19. **D-G**. Bars show percentages of total ATS that are cATS (darker bars) or iATS (lighter bars) as a function of position. Total number ATS scored: wildtype, n = 2334; 5’ Δ, n = 2305; mid Δ, n = 3086; and 3’ Δ, n = 2355. **H**. cATS frequency in wildtype control done in parallel with mutants (dashed purple line) compared to cATS frequency in wild-type from Figure 3 (solid purple line). **I-K**. cATS frequency of each mutant (solid line) compared to wildtype control (dashed purple line).

To test effects of the deletions on germline function, we assayed each 1 kb deletion mutant for fertility and PZ size. We also scored the fertility of animals homozygous for the large 5.7 kb deletion used to test intron probe specificity (large Δ). While loss of the *mpk-1b* germline isoform causes fully penetrant sterility (Robinson-Thiewes et al., in preparation), the 5’ Δ and 3’ Δ mutants were fertile and the mid Δ and large Δ mutants were mostly fertile, with a partially penetrant sterility at higher temperature (Figure 5B). Moreover, PZ length was similar to wildtype in 5’ Δ and 3’ Δ mutants and reduced by only ∼5 gcd in mid Δ and large Δ mutants (Figure 5C). We also noted that the mid Δ and large Δ mutants were vulvaless, a defect associated with MPK-1 loss in the soma (Lackner et al., 1994; Wu and Han, 1994). Therefore, mid Δ likely removes a somatic enhancer. Unexpectedly, the large Δ mutants were ∼30% dauer constitutive, a *mpk-1* defect not reported previously to our knowledge. We conclude that a 1 kb reduction in the long first *mpk-1* intron has no dramatic effect on germline function.

We also assayed transcription of each 1 kb intron deletion mutant. These deletions remove a roughly equal number of individual probes from the intron probe set (Figure 5A, Table S1); the large Δ mutant, by contrast, removes most individual probes from the intron probe set and was therefore not analyzed (Figure S1C). For each 1 kb intron deletion mutant, we performed the same experiments and analyses done with wildtype, as described above. In the distal PZ, transcriptional probability (percent cells with any ATS) was essentially the same in wildtype and the three mutants in the distal PZ (Figure S9A-H). Transcriptional output, scored either as number of nascent transcripts (Figure S9I-L) or number of mRNAs (Figure S9M-T), was similar in the three mutants and wildtype. While the gradient in percent cATS appeared to shift distally by ∼2 gcd in the mid Δ and 3’ Δ mutants (Figure 5D-G), that shift was not statistically significant (student’s t-test), and cATS frequencies were similar to the wildtype control (Figure 5H-K). Transcription in the EP region was also equivalent in wildtype and deletion mutants. We conclude that a 1 kb size reduction of the first long *mpk-1* intron (5’ Δ, mid Δ, and 3’ Δ), removal of the hairpin loop (5’ Δ), and removal of the putative somatic enhancer (mid Δ) have no significant effect on *mpk-1* transcription in the germ cells assayed.

### *mpk-1 developmental pattern of cATS* frequency *is sex-specific*

Finally, we asked if the dramatically graded increase in *mpk-1* cATS frequency that was found in the hermaphrodite PZ might reflect a developmental control related to GSC maturation. If this were the case, a similar increase would be expected in the male germline, where GSCs also reside distally and their daughters are triggered to differentiate as they leave the niche (Figure 6A) (Crittenden et al., 2019). To test this prediction, we analyzed *mpk-1* cATS frequency in adult male PZs and EPs (Figure 6A, red boxes). However, the male pattern of cATS frequency was different from that in hermaphrodites. In the distal-most GSCs within the male niche, the cATS frequency was ∼50% and increased to ∼65% by row 12 of the male PZ (Figure 6B). In the male EP region, cATS frequency dropped from ∼80% at the TZ/EP boundary to ∼40% by row 12 (Figure 6C). The *mpk-1* cATS frequency pattern is therefore sexually dimorphic (Figure 6D, 6E). The male EP decrease in cATS frequency matches well with the previously reported decrease in MPK-1 protein abundance in the same region (Min Ho Lee et al., 2007). We suggest that the sexually dimorphic patterns in cATS frequency reflect sex-specific changes in transcriptional progression that are related to sperm fate specification in the male PZ and production of maternal RNAs in the hermaphrodite EP (see Discussion).

**Figure 6:**
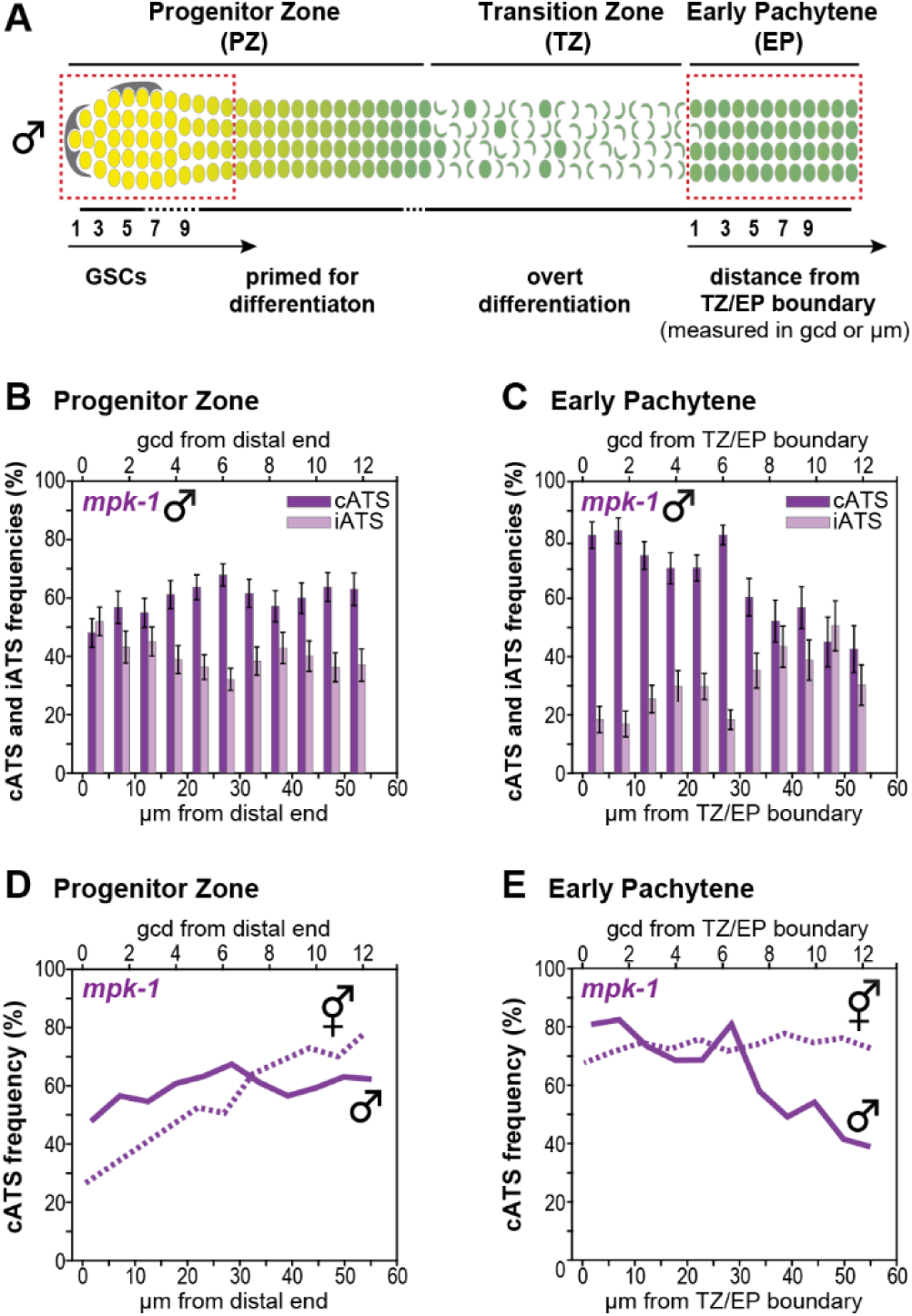
*mpk-1* ATS pattern in males. **A**. Male germline architecture. Males have two somatic niche cells (gray). Male progenitor and transition zones are longer than in hermaphrodite (Morgan et al., 2010), but male GSC pool sizes are similar in the two sexes (Crittenden et al., 2019). Red boxes, regions analyzed by smFISH, the first 12 cell rows of the PZ and the first 12 cell rows of early pachytene (EP) region. **B & D. PZ** data, conventions as in Figure 3. Number of gonads, n = 24. **C & E. EP** data, conventions as in Figure 4. Number of gonads, n = 22. **B & C**. iATS and cATS frequencies as a function of position in the PZ (B) and EP (C). Bars show percentages of total ATS that are cATS (darker bars) or iATS (lighter bars). Total number of ATS scored in PZ = 2195, and in EP = 1350. **D & E**. Male *mpk-1* cATS frequency (solid purple) compared to hermaphrodite cATS frequency (dashed purple) in the PZ (D) and EP (E). Hermaphrodite data taken from Figure 3J for PZ and Figure 4H for the EP.

## Discussion

This study analyzes transcription of three key regulatory genes during *C. elegans* germline development, using smFISH to visualize single active transcription sites (ATS) and mRNAs. Our results lead to three major conclusions. First, we identify distinct ATS classes: iATS harbor partial nascent transcripts while cATS harbor full length nascent transcripts. Second, we find that the frequencies of these ATS classes change in gene- and sex-specific fashion along the germline developmental axis, suggesting developmental regulation. Finally, we show that only one ATS class, the cATS, correlates with transcriptional productivity, suggesting an impact of ATS class on gene expression.

### Two ATS classes with distinct extents of transcriptional progression and transcriptional output

Classically, ATS are thought to harbor multiple transcripts that vary in degree of completeness (Darzacq et al., 2007; Femino et al., 1998; Mcknight and Miller, 1976). The discovery of iATS and cATS demonstrates that ATS can exist in different states with distinct extents of transcriptional progression through a gene. The iATS are detected only with the 5’ probe set and this ATS class must be dominated by partial transcripts; by contrast, cATS are detected with both 5’ and 3’ probe sets and must include complete or nearly complete transcripts, likely in addition to partial transcripts (Figure 7A). This interpretation is consistent with previous studies that used smFISH to investigate transcriptional progression (Bartman et al., 2019; Coté et al., 2020; Darzacq et al., 2007; Pichon et al., 2018). Moreover, alternative explanations seem unlikely. The lack of an overlapping exon signal at iATS might be explained if the spliced long intron was tethered to the site while full-length transcripts were released rapidly. However, introns are typically degraded quickly (Clement et al., 1999) or moved to nuclear speckles for degradation (Daguenet et al., 2012; Dias et al., 2010; Galganski et al., 2017), and a rapid release of full-length transcripts is inconsistent with the negative correlation between iATS frequency and transcriptional output (Table 1C). We therefore favor the idea that iATS and cATS sites differ in their extents of transcriptional progression (Figure 7A).

**Figure 7:**
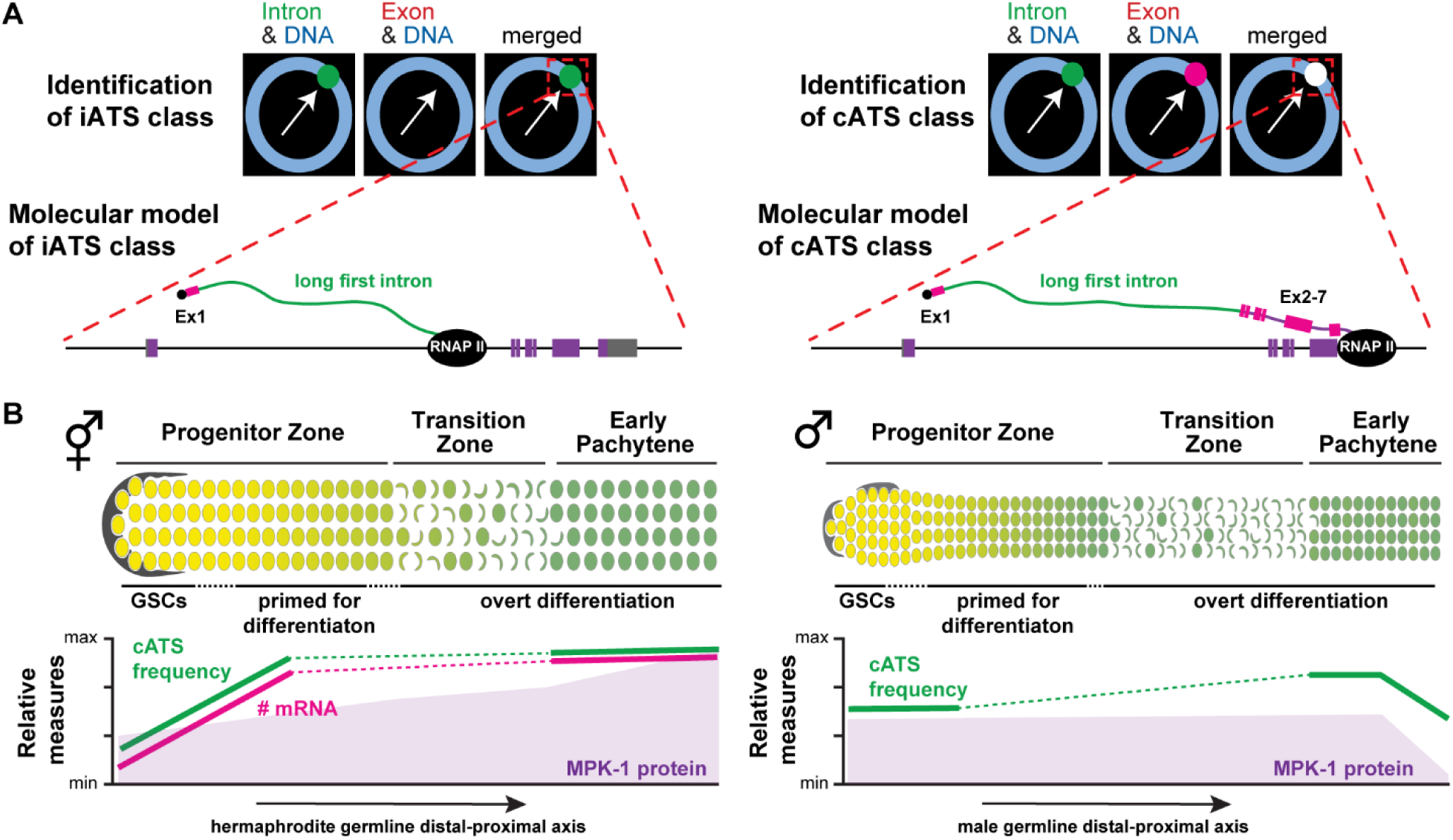
Models for ATS class regulation in development. A. Molecular model of iATS, left, and cATS, right. Above left, iATS hybridize uniquely to intron probe set; above right, cATS hybridize to both intron and exon probe sets. Below left, nascent transcripts at iATS are proposed to consist of the first exon (Ex1) plus much of the long first intron, but not more downstream exons (Ex2-7). Below right, nascent transcripts at cATS are proposed to include both the first exon, the long first intron and downstream exons. See Discussion for possible mechanisms that may be regulated to create these two ATS classes. **B**. Models for developmental effects of *mpk-1* ATS class regulation. Above, hermaphrodite (left) and male (right) gonads, using conventions as in Figure 1A. Below, patterns of *mpk-1* ATS class frequency and *mpk-1* gene expression. Patterns of cATS frequency and mRNA numbers are from this work (solid lines, dashed lines are extrapolated); patterns of MPK-1 protein abundance (purple shading) was reported earlier (Min Ho Lee et al., 2007). See Discussion for biological relevance of sexually dimorphic ATS class patterns.

We do not understand the mechanism responsible for formation of these two ATS classes. The simplest explanation is that transcriptional elongation is slowed or paused at iATS, perhaps in the long first intron. That slowing or pausing cannot be the same as “promoter-proximal pausing”, which occurs ∼20-60 bp downstream of transcriptional initiation and would not generate a transcript detectable with the first intron probe (Adelman and Lis, 2012; Jonkers and Lis, 2015; Sheridan et al., 2018). If pausing does occur at iATS, it could be coupled to splicing (Alexander et al., 2010; Mayer et al., 2015; Nojima et al., 2015; Saldi et al., 2016). However, splicing pauses detected to date in other systems are short, on the order of seconds to minutes (Alpert et al., 2017; Martin et al., 2013; Singh and Padgett, 2009). Another plausible explanation is that transcription is aborted at iATS. Abortive transcription occurs in *Drosophila* embryos, where mitosis truncates transcription of very long genes in cells with very short cell cycles (Gubb, 1986; Kwasnieski et al., 2019; Shermoen and O’Farrell, 1991; Swinburne and Silver, 2008; Tadros and Lipshitz, 2009). However, in the iATS reported in our work, elongation through each of the genes investigated is predicted to take <15 minutes, based on rates of 1-4kb/sec (Darzacq et al., 2007; Martin et al., 2013; Singh and Padgett, 2009), far less than the ∼6hr cell cycle in the progenitor zone (Albert Hubbard and Schedl, 2019); thus, iATS are unlikely to be the result of aborted transcription at mitosis. Differentiating among these various mechanisms is challenging in the *C. elegans* germline, and iATS *per se* have not yet been reported in cultured cells. Nonetheless, although speculative, we suggest that iATS result from transcriptional pausing, perhaps during RNAP II elongation or splicing of the long first intron.

### Graded cATS frequency is coupled to a graded transcriptional output

One of our most striking findings is that frequencies of the two ATS classes, iATS and cATS, are patterned in development. Whereas the probability of a nucleus harboring any ATS was essentially the same for all germ cells in the two regions investigated, the frequencies of the two ATS classes changed dramatically and reciprocally as germ cells mature. Indeed, the *mpk-1* and *lag-3* frequencies were clearly graded in the hermaphrodite progenitor zone. For both of these genes, iATS frequency started high in GSCs within the niche and dropped as their daughters left the niche and moved through the progenitor zone; conversely, cATS frequency started low and increased across the same region. These reciprocally graded iATS and cATS frequencies likely reflect graded and regulated changes in the ability of an ATS to complete transcription through the gene.

An important corollary of the iATS class with its partial transcripts is that existence of an active transcription site does not ensure production of mRNAs. The iATS frequency had a pattern opposite to that of transcriptional yield, measured both as production of full-length nascent transcripts and mature mRNAs, whereas the cATS frequency pattern aligned well with these two measures of transcriptional output. Future smFISH studies must therefore consider not only formation of ATS, but the ATS class and its productivity. This caution may be most important for genes with long introns or perhaps simply long genes. We suspected that some aspect of the long first intron might be critical for producing the two ATS classes but were not able to identify it with designer deletions. Regardless, for the genes investigated in this work, we suggest that the regulated balance between cATS and iATS classes is an important factor in driving gene expression.

### ATS classes are gene-, stage- and sex-specific during germline development

The three genes investigated in this work—*mpk-1, lag-1*, and *lag-3—*encode key regulators of development (see Introduction). In the germline tissue, they regulate stem cell maintenance, sex determination, and several steps of differentiation (Figure 1B); elsewhere, they regulate embryogenesis and post-embryonic somatic development (Priess, 2005; Sternberg, 2005). Here we consider how the patterns of ATS class frequency relate to germline function. In hermaphrodites, pachytene cells act as nurse cells for oocytes (Wolke et al., 2007) and all three genes produce maternal mRNAs (Stoeckius et al., 2014), consistent with their high cATS frequency and abundant transcriptional yield to load maternal mRNAs into the oocyte. Little is known about *lag-1* and *lag-3* function in germline differentiation; by contrast, *mpk-1* is critical (see below).

The functions of patterned cATS frequencies in the progenitor zone are more nuanced. The *lag-1* and *lag-3* genes encode essential components of the Notch-dependent transcription factor complex to maintain GSCs in response to niche signaling (Kershner et al., 2014; Lambie and Kimble, 1991; Lee et al., 2016; Petcherski and Kimble, 2000). Consistent with that role, the *lag-1* cATS frequency and transcriptional output are higher in GSCs than in the more proximal PZ (this work), and LAG-1 protein is expressed similarly (Chen et al., 2020). The *lag-3* cATS pattern was low in GSCs even though the LAG-3 protein is clearly functioning there to maintain GSCs. The distribution of LAG-3 protein is not yet known. One clue that it may be necessary to keep the LAG-3 protein a low level in GSCs is that LAG-3 forms a complex in the nucleus with the Notch intracellular domain (NICD) (Petcherski and Kimble, 2000), and abundance of nuclear NICD is vanishingly low (Crittenden et al., 1994; Sorensen et al., 2020). We suspect that LAG-3 abundance is kept low to work with its low-abundance NICD companion for Notch-dependent transcriptional activation. Moreover, mammalian Mastermind-like (MAML), a LAG-3 ortholog (Kitagawa, 2015; McElhinny et al., 2008; Wu and Griffin, 2004; Zhao et al., 2007), is often overexpressed in cancers (Forghanifard et al., 2012; Wu and Griffin, 2004). Perhaps, a low *lag-3* cATS frequency maintains a low but functional level of LAG-3 and prevents overexpression that could induce tumorigenesis.

The *mpk-1* gene encodes ERK/MAPK, which functions at several steps of germline differentiation: sperm fate specification, meiotic progression, and oocyte maturation (Arur et al., 2011; Church et al., 1995; Min Ho Lee et al., 2007; Lopez 3rd et al., 2013; Morgan et al., 2013). These functions all rely on a single germline-specific isoform, *mpk-1b* mRNA, which in hermaphrodites, generates low MPK-1B protein in the PZ and increasing MPK-1B as germ cells progress through differentiation (Min Ho Lee et al., 2007; Myon Hee Lee et al., 2007; Robinson-Thiewes et al., in preparation). As shown in Figure 7B, the pattern of *mpk-1* cATS frequency and transcriptional output in hermaphrodites conforms to the pattern of MPK-1B protein expression and its established functions in meiotic progression and oocyte maturation. Given its prominent role in germline differentiation, *mpk-1* expression might have been kept low in GSCs to maintain stem cells. However, germline sexual identity is established in the PZ (Morgan et al., 2013, 2010) and *mpk-1* also promotes sperm fate specification (Min Ho Lee et al., 2007). Indeed, the *mpk-1* cATS frequency was sexually dimorphic: 25% in GSCs of oogenic but ∼60% in GSCs of spermatogenic germline (Figure 7B). Therefore, a low cATS frequency is not required for stem cell maintenance and is consistent with a role in preventing sperm fate specification in adult hermaphrodites. More proximally, the *mpk-1* cATS frequency increases and stays high in the early pachytene region of oogenic hermaphrodites, presumably to generate its maternal load, but decreases dramatically in the early pachytene region of spermatogenic adult males. Although the abundance and activity of MPK-1/ERK are also regulated post-transcriptionally and post-translationally (Myon Hee Lee et al., 2007; Yoon et al., 2017), regulation of ATS class, and hence regulation of transcriptional progression, emerges as its earliest point of developmental control.

### Future directions

A deeper understanding of ATS classes and their regulation is a challenge for future studies. One key question that can now be addressed is whether iATS exist at other genes and outside the *C. elegans* germline –in other tissues and other species. Although this seems likely, it remains speculation for now. The literature reveals plausible iATS candidates in *Drosophila* and mouse (Shermoen and O’Farrell, 1991; Takashima et al., 2011), but the use of different methods in those studies make iATS equivalence uncertain. Thus, identifying iATS in other genes, tissues, and organisms will demonstrate their general significance and finding them in cultured cells will open their analysis to powerful biochemical and genomic methods. The *C. elegans* germline is poised to conduct more refined smFISH analyses as well as live imaging to probe the nature of iATS – does a pause occur, and if so, where in the gene and for how long? However, iATS in the *C. elegans* germline exist in limited regions, and can be of low frequency, both disadvantages for -omics methods. Regardless, the identification of a graded developmental transition from the iATS class with its partial transcripts to cATS with its complete transcripts opens the possibility of a new mode of transcriptional regulation.

## Supporting information

Supplemental images, legends, and tables

## Acknowledgements

We thank members of the Kimble lab, Wickens lab, David Brow, Bob Landick, Daniel Panaccione, and Aaron Hoskins for engaging and thoughtful discussions during the course of this work. We thank Brian Carrick, Sarah Crittenden, and Tina Lynch for critical readings of the manuscript. We also thank ChangHwan Lee for generously sharing his expertise with smFISH, MATLAB, and *lag-3* RNAi. SR was supported by the National Science Foundation Graduate Research Fellowship under grant No. (DGE-1256259) and the NIH Predoctoral training grant in Genetics 5T32GM007133. JK was an HHMI Investigator and is now supported by NIH R01 GM134119. Any opinion, findings, and conclusions or recommendations expressed in this material are those of the authors(s) and do not necessarily reflect the views of the National Science Foundation.

## Methods

### Strains and Maintenance

All strains were maintained at 20°C (Brenner, 1974) unless otherwise indicated. See Table S3 for strain names and full genotypes. Most experiments were done using wild-type N2 animals.

### Allele generation

All mutations were made using the co-CRISPR method as described (Dokshin et al., 2018; Paix et al., 2014) with guides listed in Table S4. All mutations were isolated at 25°C, but then outcrossed 2X times to wildtype and either homozygosed or balanced with qCl [qIs26] III at 20°C.

### smFISH probe design and specificity controls

All smFISH probe sets were designed using Stellaris probe designer (https://www.biosearchtech.com/stellaris-designer), using the sequence of the first intron for the intron probe set or the sequences of all exons for the exon probe set. For each probe set, a mask of 5 was used to maximize specificity. Each probe was compared to the *C. elegans* genome using BLAT (https://genome.ucsc.edu/cgi-bin/hgBlat) for independent confirmation of sequence specificity. See Table S1 for individual probe sequences and fluorophores conjugated to each probe set. RNA specificity was confirmed using α-amanitin to inhibit RNA production following a published procedure (Lee et al., 2016). Briefly, N2 mid-L4 worms were placed on a fresh plate 12 hr before the experiment. Adults were washed off the plate with M9 and incubated in 500 µL M9 with 100 µg*/*mL α-amanitin, rotating in the dark for 4 hrs. After incubation, gonads were extruded and stained using the smFISH protocol described below. *mpk-1* smFISH intron probe specificity was confirmed using the *mpk-1 (q1084)* large Δ and exon probe specificity was confirmed using the *mpk-1 (q1069)* mutation, which carries a frameshift in the *mpk-1b* specific first exon. *lag-1* and *lag-3* probe specificities were confirmed using RNAi with the protocol from (Ahringer, 2006), because mutants are either embryonic lethal, L1 lethal, or lack germline tissue (Christensen et al., 1996; Petcherski and Kimble, 2000). RNAi was performed HT115 bacteria carrying either *lag-1, lag-3*, or empty vector were grown overnight and 100 µL seeded to each plate. Bleach synchronized N2 worms were placed on empty and RNAi plates (L1s for *lag-1* and L4s for *lag-3*); for both genes, worms were grown to adulthood and their embryos processed for smFISH (see below).

### smFISH

All steps were performed under RNase free conditions: the workspace, gloves, and pipettes were routinely cleaned using RNase Zap (ThermoFisher, AM9780), RNase free tips were used, nuclease free water (ThermoFisher, AM9932) and RNase free TE (ThermoFisher, AM9849) were used to make buffers. The “original” *mpk-1* smFISH probes were used in Figure 2E and “swapped” *mpk-1* probes were used for all other experiments (Table S5).

#### Gonad smFISH

All gonads were stained at the mid L4 + 12 hr timepoint, as described (Lee et al., 2017, 2016). Briefly, mid-L4 worms were picked to plates 12 hrs before gonad dissection. Hermaphrodites or males were washed from plates with a M9+tween mixture and anaesthetized in 0.25 mM levamisole; after dissection, gonads were washed in 500 µL 1XPBS + tween (PBSTw) solution and fixed in 1 mL 3.7 % formaldehyde in PBSTw for 20 minutes, rotating at room temperature. Gonads were then washed with 1 mL PBSTw and permeabilized with 1 mL PBSTw + 0.1 % Triton-X for 10 minutes, rotating at room temperature. Gonads were then washed 2X with 1 mL PBSTw and left in 1 mL 70% ethanol (diluted in nuclease-free water) overnight at 4 °C. The next day, ethanol was replaced with a 1 mL smFISH wash (10% formamide, 2X SSC, and Tween-20) for 5 minutes. smFISH probes were diluted in hybridization buffer (10% formamide, 2X SSC, 100 mg dextran sulfate per mL of buffer) with probe concentration depending on the probe set (Table S5). Gonads were rotated at 37 °C overnight in the dark, washed in 1 mL smFISH wash + 1: 1000 DAPI (1 mg/mL) for 30 minutes, rotating in the dark at 37 °C and finally quickly washed 2X with 1 mL smFISH wash. Gonads were mounted in 18 µm Prolong Glass (ThermoFisher, P36984), cured in the dark for 2 days at room temperature, and sealed with VALAP. Slides were stored at -20 °C until imaged.

#### Embryo smFISH

Staining was modified from a previous protocol (Ji and van Oudenaarden, 2012) using smFISH solutions described above. Embryos were washed from plates with M9. After spinning and supernatant removal, bleaching solution (745 mL concentrated Clorox Bleach, 0.5 µL 4N NaOH, and 3.750 mL M9) was added, and samples rocked gently for 4 minutes at room temperature. After bleach removal, embryos were washed 3X with M9 and transferred to an RNase free microcentrifuge tube with 1000 mL of fix solution. Tubes were immediately put in liquid nitrogen to freeze crack the eggshell and then transferred to an ice bath and allowed to thaw for 20 minutes. After removal of the fix, embryos were washed 1X 1 mL PBSTw, 70% ethanol was added, and samples were left overnight at 4 °C. The next day, ethanol was replaced with 1 mL smFISH wash added for 5 minutes, which was replaced with hybridization buffer and a probe set. Hybridizing embryos were rocked gently overnight at 37 °C in the dark. The next day, embryos were washed with 1 mL smFISH + 1:1000 DAPI (1 mg/mL) and rocked again at 37 °C for 30 minutes in the dark. After DAPI, embryos were washed 2X with 1 mL smFISH wash, mounted in 18 µL of Prolong Glass and cured for 2 days in the dark at room temperature. Slides were sealed with VALAP and kept at -20 °C until imaged.

### Biological replicates

Two biological replicates were done for all experiments performed with wildtype hermaphrodites and males. To ensure the two replicates were exposed to essentially the same conditions, animals for each replicate were raised on separate OP50 plates in the same incubator; their gonads were dissected separately but in parallel; and smFISH was done separately but in parallel. For imaging, one replicate was imaged one day and the other second was imaged the next day. Each replicate was analyzed independently using the MATLAB code. For each type of experiment, the two replicates were compared for all assessed measures using the student’s t-test. For all experiments, replicates did not differ significantly and datasets were combined for presentation. For *mpk-1* replicate 1, all 20 PZ images were used in downstream analyses; 19 of 21 EP images were used for analyses. For *mpk-1* replicate 2, 17 of 20 PZ images were used; 17 of 20 EP images were used. For *lag-3* replicate 1, 13 of 20 PZ images were used for analyses; 18 of 20 EP images were used for analyses. For *lag-3* replicate 2, 19 of 20 PZ images were used; 19 of 20 EP images were used. For male *mpk-1* replicate 1, all 13 PZ images were used for analyses; 11 of 13 EP images were used for analyses. For male *mpk-1* replicate 2, 11 of 12 PZ images were used for analyses; 11 of 12 EP images were used. See MATLAB Image Processing for reasons selected images were removed. For smFISH of the mutants, wildtype was done in parallel with the mutants, but only one replicate was done—see Figure S9 legend for details.

### Image Acquisition

All smFISH stained gonads and eggs were imaged as described (Lee et al., 2016). All smFISH images were captured using a Leica SP8 confocal microscope. For TAMRA dye (651 nm), Quasar 670 (633 nm), Quasar 570 (561 nm), and far red 610 (594 nm), 2% laser power was used for excitation, while DAPI was excited using 1.5% laser power (UV 405 nm). Fluorescence signals were collected as follows: DAPI, 412-508 nm; TAMRA, 564-627 nm; Quasar 670, 650-700 nm; Quasar 570, 564-588 nm; and far red 610, 600-680 nm.

### MATLAB Image Processing

#### Threshold determination

The program RunBatch was used to determine the optimal threshold value for the relevant probe set for all images. Data from each image channel was then visualized using the program “detectionCheck_mpk1” (available on GitHub with publication). The threshold value was chosen that best captured intron and exon signals when compared to the raw image by manual inspection. Once the optimal threshold value was determined, images were further processed in MATLAB. An individual image was excluded from further analysis if the nuclear signal could not be detected and/or no threshold value could accurately represent the visible signals.

#### MATLAB image processing

MATLAB workflow is summarized in Figure S4 and based on a previous MATLAB code (Lee et al., 2016). Each channel—intron, exon, and DAPI—were independently detected. Background from the intron channel was used to generate the 3D tissue model and mark gonad boundaries in the image. Any exceptionally bright and/or irregular shaped signal outside the boundaries was discarded as a contaminant. Candidate signals were independently examined and defined from the intron and exon channels. For the intron channel, candidate signals were required to be nuclear, round, and of an intensity value greater than the background. The percent of discarded intron probe-detected signals per gonad varied: 0.5% for *mpk-1*, 0.02% for *lag-1*, and 4.3% for *lag-3*. For the exon channel, candidate nuclear signals had to be nuclear, round, and of an intensity greater than the mean intensity of a single mRNA. In addition, each signal was designated as nuclear (within DAPI boundary), in a Voronoi defined cell of 3 µm from the center of the nucleus, or in the shared cytoplasmic core or “rachis” of the germline. Exon only detected nuclear signals were discarded (see Figure S4); their detections varied per gonad depending on gene—3.2% for *mpk-1*, 3.5% for *lag-1*, and 2.8% for *lag-3*. If DAPI did not form a sphere, which happened in cells on the boundary of the image window, any nuclear ATS and cytoplasmic mRNAs were discarded.

#### MATLAB post-image analysis

Metadata from image processing were imported into MATLAB to score raw data. The intensities from intron and exon probe sets were analyzed as described (Lee et al., 2016) to confirm single molecule detection for each gene. For all three genes, over 96% of the cytoplasmic exon signals clustered around one value that was deemed to correspond to a single mRNA. For each gene, replicate 1 and replicate 2 were compared for all analyses with the student’s t-test. All data from the replicates for a single gene and condition were not statistically different and combined. Data acquired as a function of germline position were grouped into “bins” that corresponded to 5 µm windows spanning the length of the region of interest. Because cells have a 4.4 µm diameter on average, each bin approximates a row of germ cells. To calculate transcriptional probability, DAPI defined nuclei were grouped into bins by position along the germline axis. For each bin, the number of ATS positive, either iATS or cATS, cells were counted and compared to the total number of nuclei in the same bin. For number of nascent transcripts in cATS, data were again grouped in bins. For each image, the cATS signal was calculated from a comparison to the average intensity of a single cytoplasmic mRNA in the same image. cATS intensities in each bin were added together and divided by the average mRNA intensity to determine the number of nascent transcripts per cell row. To quantify mRNA abundance in cells per row, we combined cytoplasmic exon probe signals from within a cell but excluded the rachis. Binning those numbers by position generated the number of mRNAs within cells per row. To estimate the number of mRNA per cell, we divided the number of mRNA per cell row by the number of nuclei in the cell row. We calculated frequencies of iATS and cATS as their percentage of total ATS per germ cell row, rather than per cell to avoid the cases of multiple loci firing in a single cell. All ATS were divided into bins as described above and the number of iATS and cATS were counted for each bin. The counts were then converted into a percentage for each bin. Correlation tests relied on pair-wise comparisons of five “compiled” datasets (percent cells with ATS, cATS frequency, iATS frequency, number of nascent transcripts at cATS, and number of cellular mRNAs) that combined position-binned data. For example, the iATS frequency compiled dataset was made by combining the position-binned data from the iATS frequency analysis. After each compiled dataset was generated, Pearson’s correlation coefficients were calculated for pairs of datasets (see Table 1 for specific comparisons).

### Fertility assays

Worms were grown at 15°C, 20°C, or 25°C for at least one generation before scoring fertility at each temperature. Worms were bleach-synchronized and grown to adulthood (L4 + 36 hr, L4 + 24 hr, or L4 + 12 hr, respectively for each temperature). Adults were washed off plates with M9, anesthetized in 0.25 mM levamisole, mounted onto 2% agarose pads, and scored for presence of embryos using DIC on a Zeiss Axioskop microscope.

### Progenitor zone length assay

All progenitor zone lengths were scored in gonads dissected from animals raised at 20°C, as described (Crittenden et al., 2006). Briefly, gonads were dissected from L4 + 12 hr adults in 0.25 mM levamisole in PBSTw and fixed in 300 µL 2% paraformaldehyde in PBSTw for 10 minutes, rotating at room temperature. Gonads were then washed in 1 mL PBSTw 1X, permeabilized in 1 mL PBSTw + 0.5% BSA + 0.1% Triton-X for 10 minutes, rotating at room temperature, incubated with 1:1000 DAPI (1 mg/mL) for 30 minutes, rotating in the dark at room temperature, and washed 3X in 1 mL PBSTw. After removing excess liquid, gonads were mounted in 10 *µL* Vectashield and kept at 4°C until imaged using the 63/1.4 NA Plan Apochromat oil immersion objective of a Zeiss Axioskop microscope. DAPI was visualized using the Carl Zeiss filter set 49. Images were taken as previous described (Haupt et al., 2019).

### Statistical analyses

All statistical tests were performed in MATLAB: student’s t-test (ttest2 function) and Pearson’s correlation tests (corr function). Significance cutoff of p ≤ 0.01 was used.

## Notes

### Competing Interest Statement

The authors have declared no competing interest.

## References

Adelman K, Lis JT. 2012. Promoter-proximal pausing of RNA polymerase II: emerging roles in metazoans. Nat Rev Genet. doi:10.1038/nrg3293

Ahringer J (ed.. 2006. Reverse genetics. WormBook. doi:10.1895/wormbook.1.47.1

Albert Hubbard EJ, Schedl T. 2019. Biology of the caenorhabditis elegans germline stem cell system. Genetics 213:1145–1188. doi:10.1534/genetics.119.300238

Alexander RD, Innocente SA, Barrass JD, Beggs JD. 2010. Splicing-Dependent RNA polymerase pausing in yeast. Mol Cell 40:582–593. doi:10.1016/j.molcel.2010.11.005

Alpert T, Herzel L, Neugebauer KM. 2017. Perfect timing: splicing and transcription rates in living cells. Wiley Interdiscip Rev RNA. doi:10.1002/wrna.1401

Arur S, Ohmachi M, Berkseth M, Nayak S, Hansen D, Zarkower D, Schedl T. 2011. MPK-1 ERK controls membrane organization in C. elegans oogenesis via a sex-determination module. Dev Cell 20:677–688. doi:10.1016/j.devcel.2011.04.009

Bartman CR, Hamagami N, Keller CA, Giardine B, Hardison RC, Blobel GA, Raj A. 2019. Transcriptional Burst Initiation and Polymerase Pause Release Are Key Control Points of Transcriptional Regulation. Mol Cell 73:519–532.e4. doi:10.1016/j.molcel.2018.11.004

Bender TP, Thompson CB, Kuehl WM. 1987. Differential eression of c-myb mRNA in mrine B lymphomas by a block to transcription elongation. Science (80-) 237:1473–1476. doi:10.1126/science.3498214

Brenner S. 1974. The genetics of Caenorhabditis elegans. Genetics 77:71–94.

Chen J, Mohammad A, Pazdernik N, Huang H, Bowman B, Tycksen E, Schedl T. 2020. GLP-1 Notch—LAG-1 CSL control of the germline stem cell fate is mediated by transcriptional targets lst-1 and sygl-1. PLOS Genet 16:e1008650. doi:10.1371/journal.pgen.1008650

Chorev M, Carmel L. 2012. The Function of Introns. Front Genet 3:55. doi:10.3389/fgene.2012.00055

Christensen S, Kodoyianni V, Bosenberg M, Friedman L, Kimble J. 1996. lag-1, a gene required for lin-12 and glp-1 signaling in Caenorhabditis elegans, is homologous to human CBF1 and Drosophila Su(H). Development 122:1373–1383.

Chubb JR, Trcek T, Shenoy SM, Singer RH. 2006. Transcriptional pulsing of a developmental gene. Curr Biol 16:1018–1025. doi:10.1016/j.cub.2006.03.092

Church DL, Guan KL, Lambie EJ. 1995. Three genes of the MAP kinase cascade, mek-2, mpk-1/sur-1 and let-60 ras, are required for meiotic cell cycle progression in Caenorhabditis elegans. Development 121:2525–2535.

Churchman LS, Weissman JS. 2011. Nascent transcript sequencing visualizes transcription at nucleotide resolution. Nature 469:368–373. doi:10.1038/nature09652

Cinghu S, Yang P, Kosak JP, Conway AE, Kumar D, Oldfield AJ, Adelman K, Jothi R. 2017. Intragenic Enhancers Attenuate Host Gene Expression. Mol Cell 68:104-117.e6. doi:10.1016/j.molcel.2017.09.010

Cinquin O, Crittenden SL, Morgan DE, Kimble J. 2010. Progression from a stem cell-like state to early differentiation in the C. elegans germ line. Proc Natl Acad Sci U S A 107:2048–2053. doi:10.1073/pnas.0912704107

Clement JQ, Qian L, Kaplinsky N, Wilkinson MF. 1999. The stability and fate of a spliced intron from vertebrate cells. RNA 5:206–220. doi:10.1017/S1355838299981190

Coté A, Coté C, Bayatpour S, Drexler HL, Alexander KA, Chen F, Wassie AT, Boyden ES, Berger S, Churchman LS, Raj A. 2020. The spatial distributions of pre-mRNAs suggest post-transcriptional splicing of specific introns within endogenous genes. bioRxiv 2020.04.06.028092. doi:10.1101/2020.04.06.028092

Crittenden SL, Lee C, Mohanty I, Battula S, Knobel K, Kimble J. 2019. Sexual dimorphism of niche architecture and regulation of the Caenorhabditis elegans germline stem cell pool. Mol Biol Cell 30:1757–1769. doi:10.1091/mbc.E19-03-0164

Crittenden SL, Leonhard KA, Byrd DT, Kimble J. 2006. Cellular analyses of the mitotic region in the Caenorhabditis elegans adult germ line. Mol Biol Cell 17:3051–3061. doi:10.1091/mbc.E06-03-0170

Crittenden SL, Troemel ER, Evans TC, Kimble J. 1994. GLP-1 is localized to the mitotic region of the C. elegans germ line. Development 120:2901–2911.

Daguenet E, Baguet A, Degot S, Schmidt U, Alpy F, Wendling C, Spiegelhalter C, Kessler P, Rio MC, Hir H Le, Bertrand E, Tomasetto C. 2012. Perispeckles are major assembly sites for the exon junction core complex. Mol Biol Cell 23:1765–1782. doi:10.1091/mbc.E12-01-0040

Darzacq X, Shav-Tal Y, De Turris V, Brody Y, Shenoy SM, Phair RD, Singer RH. 2007. In vivo dynamics of RNA polymerase II transcription. Nat Struct Mol Biol 14:796–806. doi:10.1038/nsmb1280

Dias AP, Dufu K, Lei H, Reed R. 2010. A role for TREX components in the release of spliced mRNA from nuclear speckle domains. Nat Commun 1:1–10. doi:10.1038/ncomms1103

Dokshin GA, Ghanta KS, Piscopo KM, Mello CC. 2018. Robust Genome Editing With Short Single-Stranded and Long, Partially Single-Stranded DNA Donors in Caenorhabditiselegans. Genetics. doi:10.1534/genetics.118.301532

Femino AM, Fay FS, Fogarty K, Singer RH. 1998. Visualization of single RNA transcripts in situ. Science (80-) 280:585–590. doi:10.1126/science.280.5363.585

Forghanifard MM, Moaven O, Farshchian M, Montazer M, Raeisossadati R, Abdollahi A, Moghbeli M, Nejadsattari T, Parivar K, Abbaszadegan MR. 2012. Expression analysis elucidates the roles of MAML1 and Twist1 in esophageal squamous cell carcinoma aggressiveness and metastasis. Ann Surg Oncol 19:743–749. doi:10.1245/s10434-011-2074-8

Galganski L, Urbanek MO, Krzyzosiak WJ. 2017. SURVEY AND SUMMARY Nuclear speckles: molecular organization, biological function and role in disease. Nucleic Acids Res 45:10350–10368. doi:10.1093/nar/gkx759

Gubb D. 1986. Intron-delay and the precision of expression of homoeotic gene products in Drosophila. Dev Genet 7:119–131. doi:10.1002/dvg.1020070302

Haupt KA, Law KT, Enright AL, Kanzler CR, Shin H, Wickens M, Kimble J. 2019. A PUF Hub Drives Self-Renewal in Caenorhabditis elegans Germline Stem Cells. Genetics. doi:10.1534/genetics.119.302772

Ji N, van Oudenaarden A. 2012. Single molecule fluorescent in situ hybridization (smFISH) of C. elegans worms and embryos. WormBook. doi:10.1895/wormbook.1.153.1

Jonkers I, Lis JT. 2015. Getting up to speed with transcription elongation by RNA polymerase II. Nat Rev Mol Cell Biol. doi:10.1038/nrm3953

Kershner AM, Shin H, Hansen TJ, Kimble J. 2014. Discovery of two GLP-1/Notch target genes that account for the role of GLP-1/Notch signaling in stem cell maintenance. Proc Natl Acad Sci U S A 111:3739–3744. doi:10.1073/pnas.1401861111

Kitagawa M. 2015. Notch signalling in the nucleus: Roles of Mastermind-like (MAML) transcriptional coactivators. J Biochem. doi:10.1093/jb/mvv123

Kwasnieski JC, Orr-Weaver TL, Bartel DP. 2019. Early genome activation in Drosophila is extensive with an initial tendency for aborted transcripts and retained introns. Genome Res 29:1188–1197. doi:10.1101/gr.242164.118

Lackner MR, Kornfeld K, Miller LM, Robert Horvitz H, Kim SK. 1994. A MAP kinase homolog, mpk-1, is involved in ras-mediated induction of vulval cell fates in Caenorhabditis elegans. Genes Dev 8:160–173. doi:10.1101/gad.8.2.160

Lambie EJ, Kimble J. 1991. Two homologous regulatory genes, lin-12 and glp-1, have overlapping functions. Development 112:231–240.

Lee C, Seidel HS, Lynch TR, Sorensen EB, Crittenden SL, Kimble J. 2017. Single-molecule RNA fluorescence in situ hybridization (smFISH) in Caenorhabditis elegans. Bio-protocol 7:e2357. doi:10.21769/BioProtoc.2357

Lee C, Shin H, Kimble J. 2019. Dynamics of Notch-dependent transcriptional bursting in its native context. Dev Cell in press. doi:10.1016/j.devcel.2019.07.001

Lee C, Sorensen EB, Lynch TR, Kimble J. 2016. C. elegans GLP-1/Notch activates transcription in a probability gradient across the germline stem cell pool. Elife 5:e18370. doi:10.7554/elife.18370

Lee Myon Hee, Hook B, Pan G, Kershner AM, Merritt C, Seydoux G, Thomson JA, Wickens M, Kimble J. 2007. Conserved regulation of MAP kinase expression by PUF RNA-binding proteins. PLoS Genet 3:2540–2550. doi:10.1371/journal.pgen.0030233

Lee Min Ho, Ohmachi M, Arur S, Nayak S, Francis R, Church D, Lambie E, Schedl T. 2007. Multiple functions and dynamic activation of MPK-1 extracellular signal-regulated kinase signaling in Caenorhabditis elegans germline development. Genetics 177:2039–2062. doi:10.1534/genetics.107.081356

Liu T, Rechtsteiner A, Egelhofer TA, Vielle A, Latorre I, Cheung MS, Ercan S, Ikegami K, Jensen M, Kolasinska-Zwierz P, Rosenbaum H, Shin H, Taing S, Takasaki T, Iniguez AL, Desai A, Dernburg AF, Kimura H, Lieb JD, Ahringer J, Strome S, Liu XS. 2011. Broad chromosomal domains of histone modification patterns in C. elegans. Genome Res 21:227–236. doi:10.1101/gr.115519.110

Lopez 3rd AL, Chen J, Joo HJ, Drake M, Shidate M, Kseib C, Arur S. 2013. DAF-2 and ERK couple nutrient availability to meiotic progression during Caenorhabditis elegans oogenesis. Dev Cell 27:227–240. doi:10.1016/j.devcel.2013.09.008

Mannervik M, Nibu Y, Zhang H, Levine M. 1999. Transcriptional coregulators in development. Science (80-) 284:606–609.

Martin RM, Rino J, Carvalho C, Kirchhausen T, Carmo-Fonseca M. 2013. Live-Cell Visualization of Pre-mRNA Splicing with Single-Molecule Sensitivity. Cell Rep 4:1144–1155. doi:10.1016/j.celrep.2013.08.013

Mayer A, Di Iulio J, Maleri S, Eser U, Vierstra J, Reynolds A, Sandstrom R, Stamatoyannopoulos JA, Churchman LS. 2015. Native elongating transcript sequencing reveals human transcriptional activity at nucleotide resolution. Cell 161:541–554. doi:10.1016/j.cell.2015.03.010

McElhinny AS, Li JL, Wu L. 2008. Mastermind-like transcriptional co-activators: Emerging roles in regulating cross talk among multiple signaling pathways. Oncogene. doi:10.1038/onc.2008.228

Mcknight SL, Miller OL. 1976. Ultrastructural Patterns of RNA Synthesis during Early Embryogenesis of Drosophila melanogaster, Cell.

Morgan CT, Noble D, Kimble J. 2013. Mitosis-meiosis and sperm-oocyte fate decisions are separable regulatory events. Proc Natl Acad Sci U S A 110:3411–3416. doi:10.1073/pnas.1300928110

Morgan DE, Crittenden SL, Kimble J. 2010. The C. elegans adult male germline: Stem cells and sexual dimorphism. Dev Biol 346:204–214. doi:10.1016/j.ydbio.2010.07.022

Nojima T, Gomes T, Grosso ARF, Kimura H, Dye MJ, Dhir S, Carmo-Fonseca M, Proudfoot NJ. 2015. Mammalian NET-seq reveals genome-wide nascent transcription coupled to RNA processing. Cell 161:526–540. doi:10.1016/j.cell.2015.03.027

Paix A, Wang Y, Smith HE, Lee CY, Calidas D, Lu T, Smith J, Schmidt H, Krause MW, Seydoux G. 2014. Scalable and versatile genome editing using linear DNAs with microhomology to Cas9 Sites in Caenorhabditis elegans. Genetics 198:1347–1356. doi:10.1534/genetics.114.170423

Parenteau J, Maignon L, Berthoumieux M, Catala M, Gagnon V, Abou Elela S. 2019. Introns are mediators of cell response to starvation. Nature 565:612–617. doi:10.1038/s41586-018-0859-7

Pereira LA, Hugo HJ, Malaterre J, Huiling X, Sonza S, Cures A, J Purcell DF, Ramsland PA, Gerondakis S, Gonda TJ, Ramsay RG. 2015. MYB Elongation Is Regulated by the Nucleic Acid Binding of NFκB p50 to the Intronic Stem-Loop Region. doi:10.1371/journal.pone.0122919

Petcherski AG, Kimble J. 2000. LAG-3 is a putative transcriptional activator in the C. elegans Notch pathway. Nature 405:364–368. doi:10.1038/35012645

Pichon X, Lagha M, Mueller F, Bertrand E. 2018. A Growing Toolbox to Image Gene Expression in Single Cells: Sensitive Approaches for Demanding Challenges. Mol Cell. doi:10.1016/j.molcel.2018.07.022

Priess JR. 2005. Notch signaling in the C. elegans embryo. WormBook. doi:10.1895/wormbook.1.4.1

Rose AB. 2019. Introns as gene regulators: A brick on the accelerator. Front Genet. doi:10.3389/fgene.2018.00672

Rosu S, Cohen-Fix O. 2017. Live-imaging analysis of germ cell proliferation in the C. elegans adult supports a stochastic model for stem cell proliferation. Dev Biol 423:93–100. doi:10.1016/j.ydbio.2017.02.008

Saldi T, Cortazar MA, Sheridan RM, Bentley DL. 2016. Coupling of RNA Polymerase II Transcription Elongation with Pre-mRNA Splicing. J Mol Biol. doi:10.1016/j.jmb.2016.04.017

Sheridan RM, Fong N, D’Alessandro A, Bentley DL. 2018. Widespread Backtracking by RNA Pol II Is a Major Effector of Gene Activation, 5′ Pause Release, Termination, and Transcription Elongation Rate. Mol Cell. doi:10.1016/j.molcel.2018.10.031

Shermoen AW, O’Farrell PH. 1991. Progression of the cell cycle through mitosis leads to abortion of nascent transcripts. Cell 67:303–310. doi:10.1016/0092-8674(91)90182-x

Singh J, Padgett RA. 2009. Rates of in situ transcription and splicing in large human genes. Nat Struct Mol Biol 16:1128–1133. doi:10.1038/nsmb.1666

Sorensen EB, Seidel HS, Crittenden SL, Ballard JH, Kimble J. 2020. A toolkit of tagged glp-1 alleles reveals strong glp-1 expression in the germline, embryo, and spermatheca.

Sternberg PW. 2005. Vulval development. WormBook. doi:doi/10.1895/wormbook.1.6.1

Stoeckius M, Grün D, Kirchner M, Ayoub S, Torti F, Piano F, Herzog M, Selbach M, Rajewsky N. 2014. Global characterization of the oocyte-to-embryo transition in C aenorhabditis elegans uncovers a novel m RNA clearance mechanism. EMBO J 33:1751–1766. doi:10.15252/embj.201488769

Swinburne IA, Miguez DG, Landgraf D, Silver PA. 2008. Intron length increases oscillatory periods of gene expression in animal cells. Genes Dev 22:2342–2346. doi:10.1101/gad.1696108

Swinburne IA, Silver PA. 2008. Intron Delays and Transcriptional Timing during Development. Dev Cell 14:324–330. doi:10.1016/j.devcel.2008.02.002

Tadros W, Lipshitz HD. 2009. The maternal-to-zygotic transition: A play in two acts. Development 136:3033–3042. doi:10.1242/dev.033183

Takashima Y, Ohtsuka T, González A, Miyachi H, Kageyama R. 2011. Intronic delay is essential for oscillatory expression in the segmentation clock. Proc Natl Acad Sci U S A 108:3300–3305. doi:10.1073/pnas.1014418108

Wolke U, Jezuit EA, Priess JR. 2007. Actin-dependent cytoplasmic streaming in C. elegans oogenesis. Development 134:2227–2236. doi:10.1242/dev.004952

Wu L, Griffin JD. 2004. Modulation of Notch signaling by mastermind-like (MAML) transcriptional co-activators and their involvement in tumorigenesis. Semin Cancer Biol 14:348–356. doi:10.1016/j.semcancer.2004.04.014

Wu Y, Han M. 1994. Suppression of activated let-60 ras protein defines a role of Caenorhabditis elegans sur-1 MAP kinase in vulval differentiation. Genes Dev 8:147–159. doi:10.1101/gad.8.2.147

Yoon DS, Alfhili MA, Friend K, Lee M-H. 2017. MPK-1/ERK regulatory network controls the number of sperm by regulating timing of sperm-oocyte switch in C. elegans germline. Biochem Biophys Res Commun 491:1077–1082. doi:10.1016/j.bbrc.2017.08.014

Zhao Y, Katzman RB, Delmolino LM, Bhat I, Zhang Y, Gurumurthy CB, Germaniuk-Kurowska A, Reddi H V., Solomon A, Zeng M-S, Kung A, Ma H, Gao Q, Dimri G, Stanculescu A, Miele L, Wu L, Griffin JD, Wazer DE, Band H, Band V. 2007. The Notch Regulator MAML1 Interacts with p53 and Functions as a Coactivator. J Biol Chem 282:11969–11981. doi:10.1074/jbc.M608974200

