## Supplemental images, legends, and tables for "Two classes of active transcription sites and their roles in developmental regulation"

Supplemental Figures:

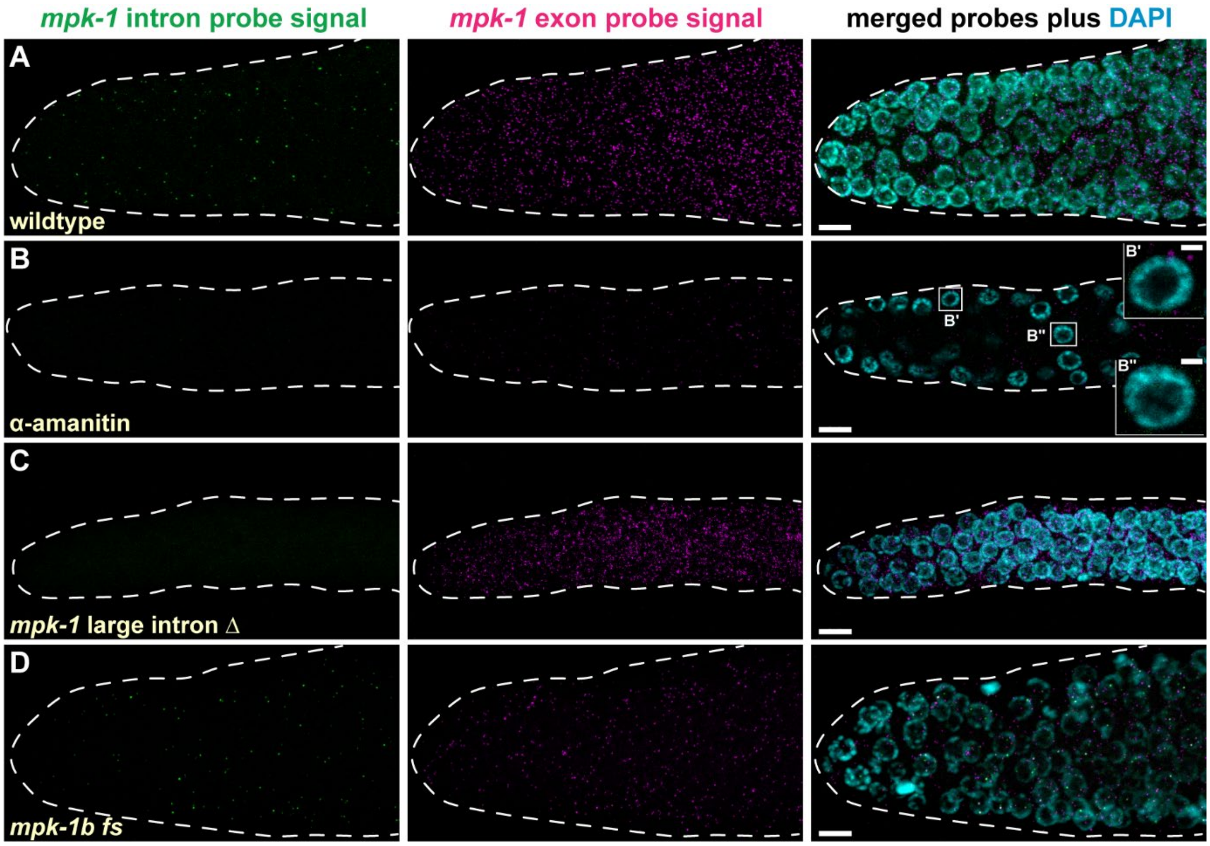


**Figure S1: *mpk-1* smFISH specificity controls. A-D.** Representative dissected gonads stained for *mpk-1* RNA. The experimental condition (e.g. wildtype) noted in the first image of each row applies to all images in that row. Each column represents signal from the intron probe set (green), signal from the exon probe set (magenta), or a composite of the two probe sets with DAPI (cyan). Scale, 5 µm. **A.** Wildtype progenitor zone shown as a maximum projection; image is same as in Figure 2A. **B** Progenitor zone after α-amanitin treatment and shown as a single z-plane. Treatment removes most nuclear signal seen either with intron or exon probes. **B'-B''.** Insets are representative magnified nuclei that lack ATS. Scale, 1 µm. **C.** *mpk-1* mutant gonad homozygous for a large deletion removing most intron probe binding sites (see Figure 1C for deletion size and position). Intron probe signal is eliminated, but cytoplasmic exon probe signal remains. Maximum projection shown. **D.** *mpk-1* mutant gonad homozygous for a frameshift (fs)-causing 1 bp insertion in the first *mpk-1b* exon that creates a premature stop codon in the germline-specific *mpk-1b* isoform. Intron probe signals remain, but exon probe signal is largely reduced (compare panel A to D, exon probe signal). Maximum projection shown.

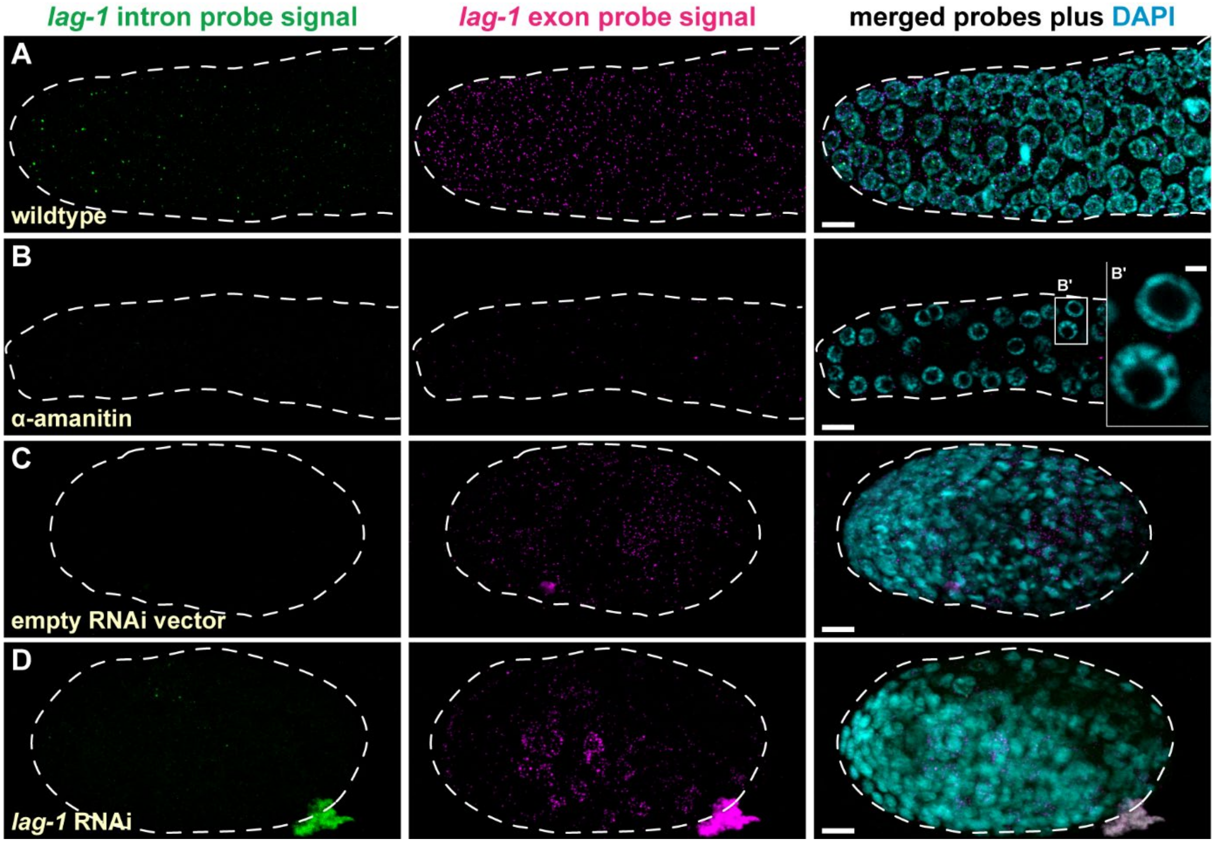


**Figure S2: *lag-1* smFISH specificity controls. A-D.** Representative dissected gonads or embryos stained for *lag-1* RNA. Conventions are same as Figure S1. Scale, 5 µm. **A.** Wildtype progenitor zone shown as a maximum projection; image is same as in Figure 2B. **B.** Progenitor zone stained after α-amanitin treatment and shown as a single z-plane. Treatment removes most nuclear signal seen either with intron or exon probes. **B'**. Magnified representative nuclei confirm lack of ATS. Scale, 1 µm. **C.** Embryo stained after treatment with empty vector RNAi as a control and shown as a maximum projection. An embryo is shown for comparison with *lag-1* RNAi, which causes L1 larval lethality. **D.** Embryo stained after *lag-1* RNAi and shown as a maximum projection. *lag-1* mRNA changes from diffuse cytoplasmic dots to perinuclear granules. Although not an expected effect of RNAi, *lag-1* RNAi leads to a Lag-1 phenotype as expected after *lag-1* depletion; a similar change in RNA localization occurs with RNAi directed against another gene, again with gene-specific phenotypic defects (S. Crittenden and J. Kimble, unpublished).

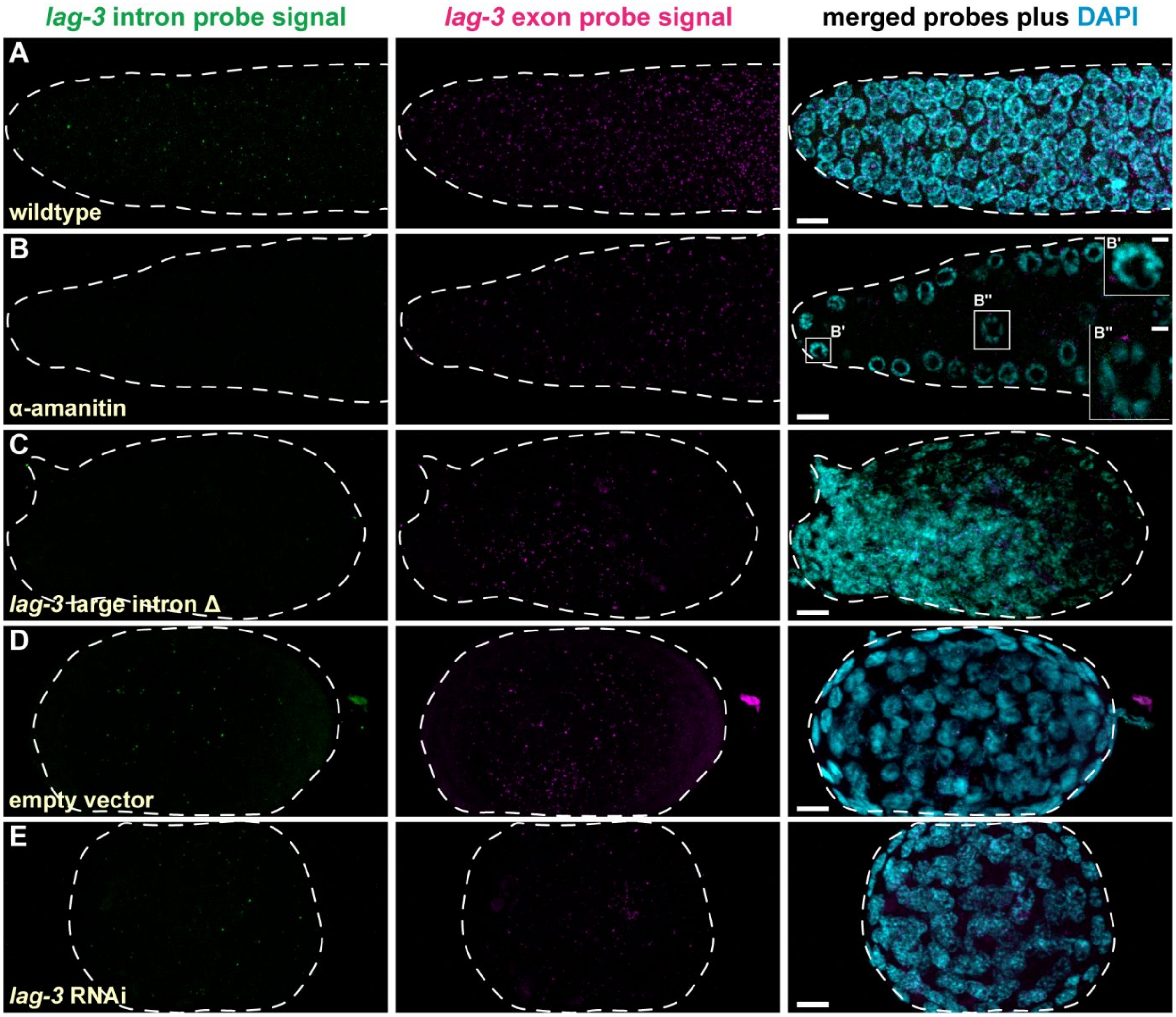


**Figure S3: *lag-3* smFISH specificity controls. A-D.** Representative dissected gonads or embryos stained for *lag-3* RNA. Conventions are same as Figure S1. Scale, 5 µm. **A.** Wildtype progenitor zone shown as a maximum projection; image is same as in Figure 2C. **B.** Progenitor zone stained after α-amanitin treatment and shown as a single z-plane. Treatment removes most nuclear signal seen either with intron or exon probes. **B'-B''**. Magnified nuclei confirm lack of ATS. Scale, 1 µm. **C.** *lag-3* mutant embryo homozygous for a large deletion that removes most intron probe binding sites (see Figure 1E for deletion size and position). This deletion is embryonic lethal. Maximum projection shown. **D.** Embryo stained after treatment with empty vector RNAi as a control. An embryo is shown for comparison with *lag-3* RNAi, as *lag-3* RNAi treatment causes embryonic lethality. Maximum projection shown. **E.** Embryo stained after *lag-3* RNAi. *lag-3* RNAi reduces the abundance of *lag-3* mRNA, which is the expected RNAi effect.

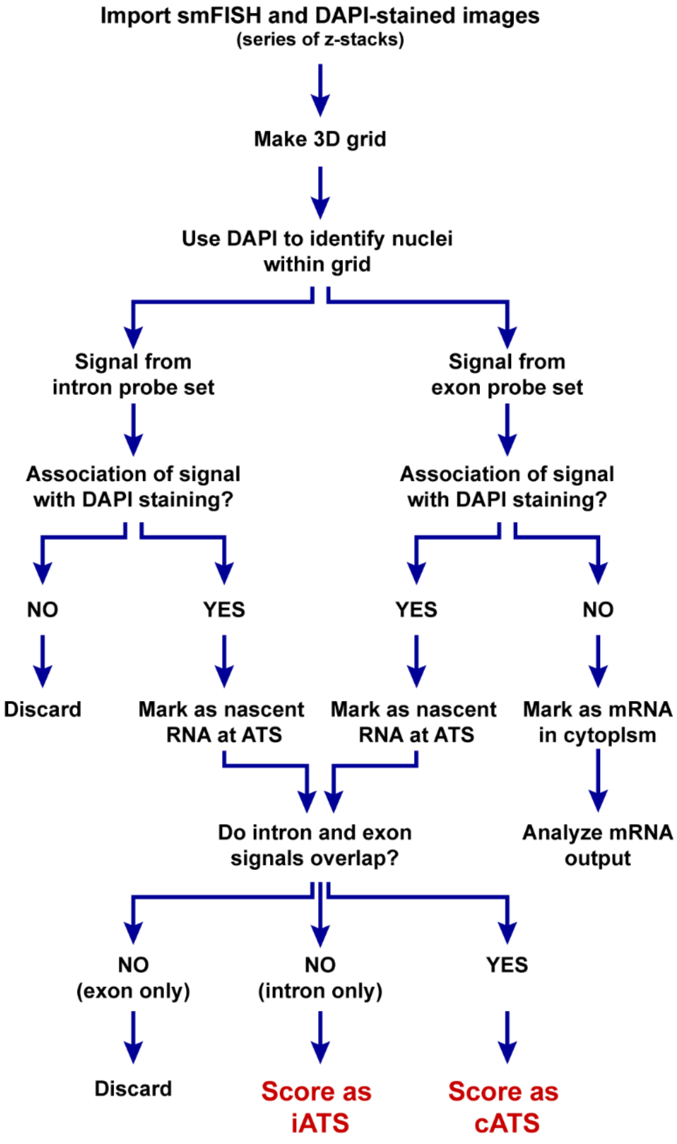


**Figure S4: MATLAB detection of iATS and ATS.** Images were imported into MATLAB for analysis. For each image, a 3D grid of the gonad was created using the background signal from the intron channel. Next, the DAPI channel was used to identify and place nuclei within the grid. Signals from the intron and exon channels were analyzed independently to generate a candidate list of intron or exon signals. The candidate lists were then independently tested for association with DAPI stained DNA. To be scored as a potential active transcription site (ATS), a candidate signal was required to overlap with DAPI and be circular (Lee et al., 2016). Intron signals that overlapped with DAPI were scored as ATS and those not overlapping were discarded. The percentages intron probe signals outside the nucleus varied: *mpk-1,* 0.5%; *lag-1,* 0.02%; and *lag-3*, 4.3%. Exon signals that overlapped with DAPI were scored as nuclear and those that did not overlap were scored as cytoplasmic (Lee et al., 2016). Next the resultant nuclear intron-detected and exon-detected signals were compared to each other for overlap within the same nucleus. If intron and exon signals were within 1.5 µm of each other in the x, y, and z direction, they were scored as a “complete” active transcription site (cATS). If intron signals did not overlap with an exon signal, they were scored as an “incomplete” active transcription site (iATS). By visual inspection, exon-only signals in the nucleus were rare and might be explained as either an exon-specific ATS or an mRNA in transport from the nucleus; thus, they were discarded. The percentages of “exon-only” nuclear signals in the nucleus were the following: *mpk-1,* 3.2%; *lag-1,* 3.5%; and *lag-3,* 2.8%.

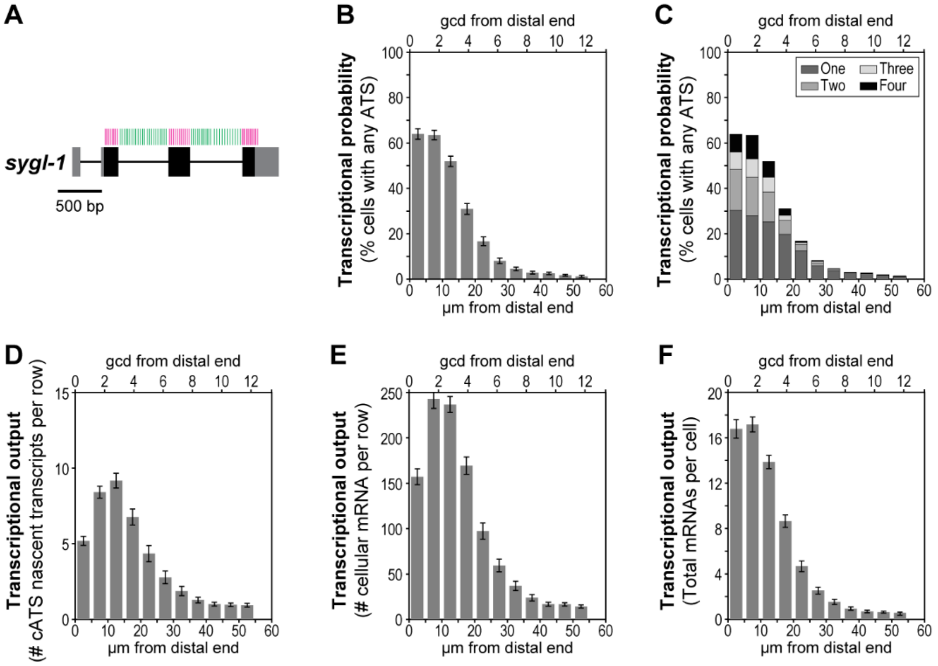


**Figure S5: MATLAB code control.** The modified MATLAB code used in this study was tested against published *sygl-1* smFISH images that had been analyzed with the original code (Lee et al., 2016). Data in B, C and F are directly comparable to published values for the image set (Lee et al., 2016). Data in D and E are additional analyses to show results obtained for *sygl-1* with the new code. **A.** *sygl-1* gene showing sites of individual smFISH probes in the exon probe set (magenta) and intron probe set (green). Conventions are same as in Figure 1C-E. Note that intron and exon probe sets are evenly distributed across the gene body so they cannot be used to assess transcriptional progression. **B-F.** smFISH analyses as a function of position in the progenitor zone. x-axis, positions measured in gcd (top) and µm (bottom) from the distal end. Standard error shown for each bar graph. **B.** Transcriptional probability for *sygl-1,* measured as percentage of cells with at least one ATS, including iATS and cATS. Compare to Figure 3A in (Lee et al., 2016). **C.** Transcriptional probability broken down by proportion with number of ATS per nucleus, from one to four (see legend top right). Compare to Figure 3C in (Lee et al., 2016). **D.** Number of nascent transcripts at cATS per cell row, calculated as described in Figure 3 legend and methods. **E.** The number of *sygl-1* mRNA in cells (not rachis), per row. **F.** Number of *sygl-1* mRNA per cell. Compare to Figure 3G in (Lee et al., 2016).

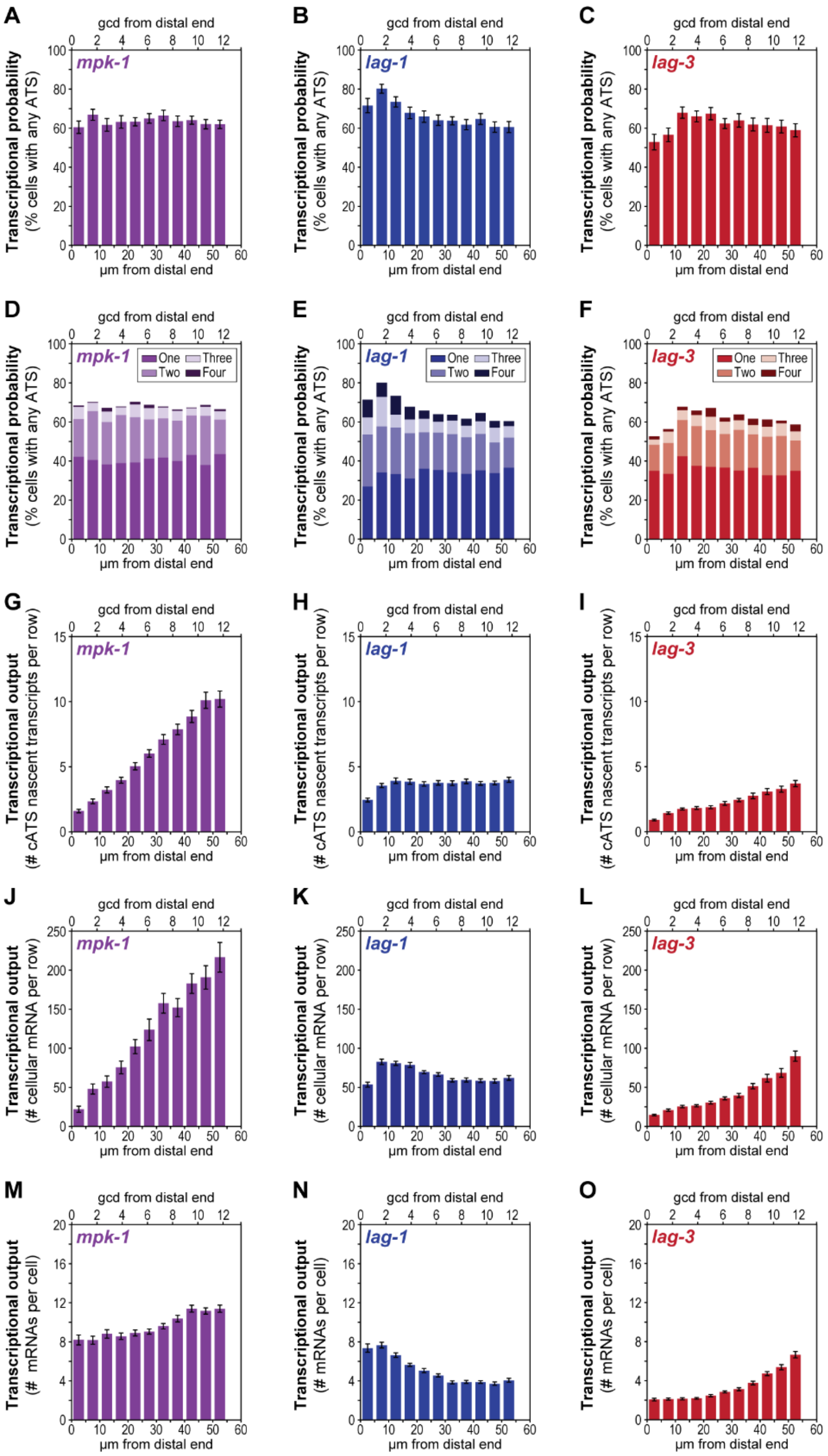


**Figure S6: Expanded data for Figure 3**. Transcription analyses in distal progenitor zone (dashed red box in Figure 3A). x-axis, gcd (top) and µm (bottom) from the distal end. Standard errors are shown for each bar graph. **A-C.** Transcriptional probability measured as percentage of cells with at least one ATS, including iATS and cATS. **D-F**. Transcriptional probability broken down by proportion with ATS numbers from one to four (see legend top right). **G-I.** Transcriptional output measured as total number of nascent transcripts at cATS per cell row. **J-L.** Transcriptional output measured as total number of cellular mRNAs (not rachis) per cell row. Germ cell boundaries were determined from MATLAB-generated Voronoi cells centered in the nucleus (see Methods). **M-O.** Transcriptional output measured as number of mRNAs within each cell (not rachis) as a function of position.

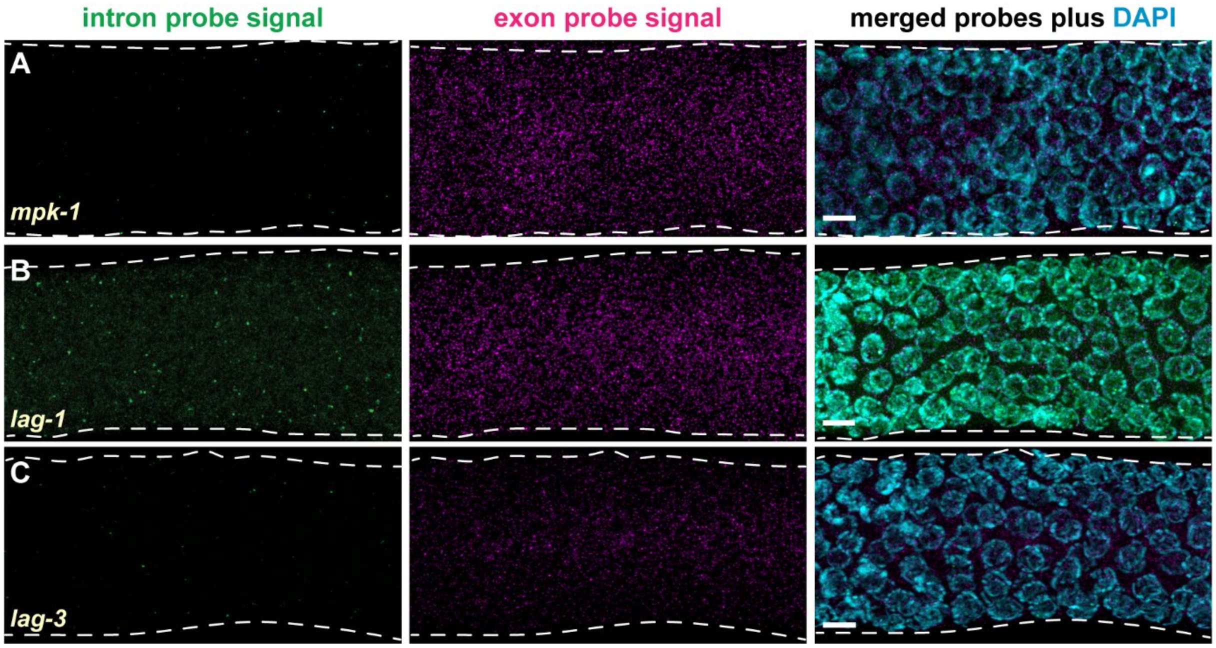


**Figure S7: Representative smFISH images in early pachytene region. A-C.** Images are arranged as in Figure 2A-C. Each row shows results from a different gene; each column shows results from a different probe signal, either alone – left, intron (green), middle, exon (magenta) – or merged (right) with DAPI (cyan). Scale, 5 µm.

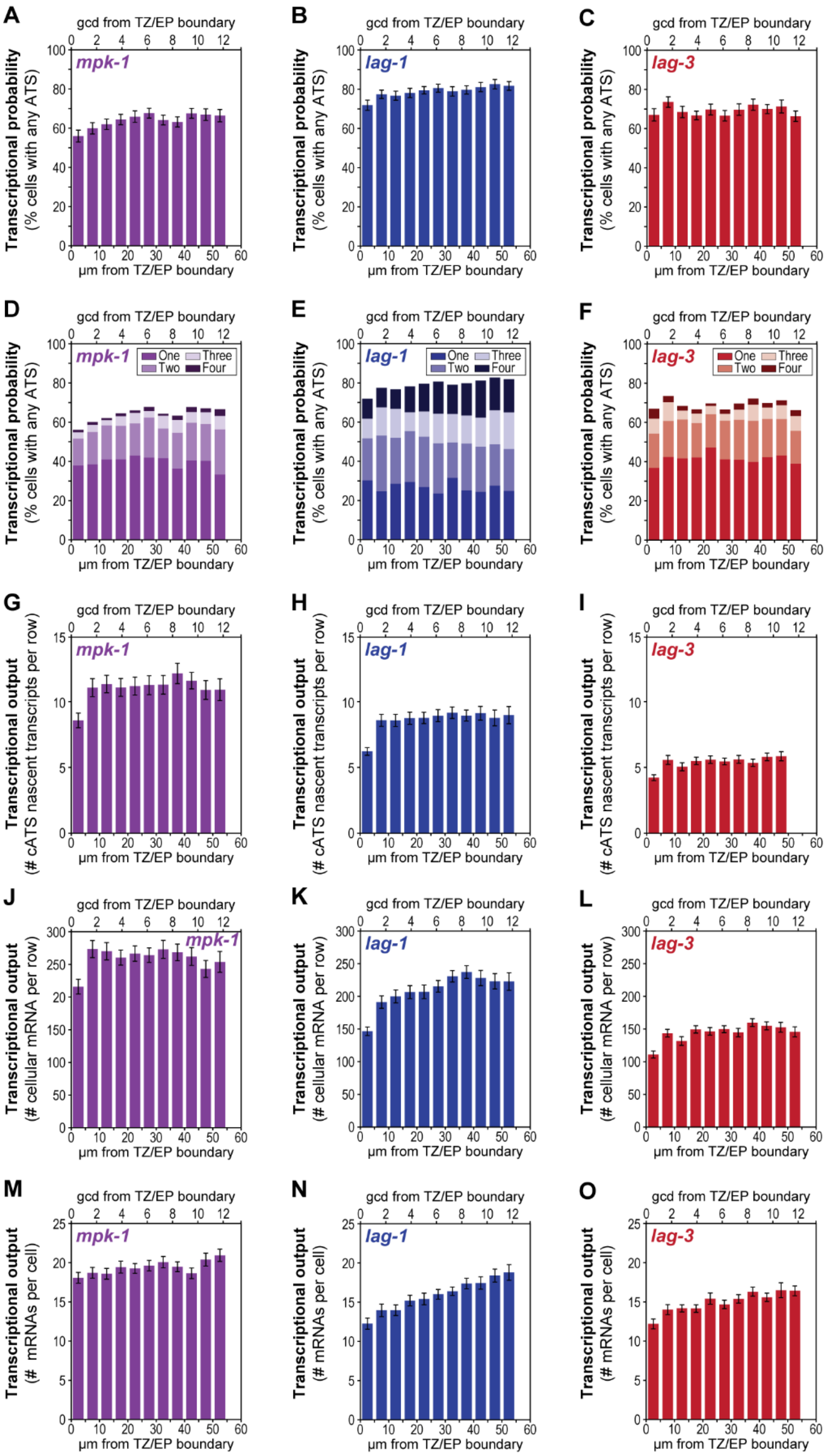


**Figure S8: Expanded data for Figure 4.** Transcription analyses in the early pachytene region (dashed red box in Figure 4A). x-axis, gcd (top) and μm (bottom) from the boundary between the Transition Zone (TZ) and Early Pachytene (EP) region. Standard errors are shown for each bar graph. **A-C.** Transcriptional probability measured as percentage of cells with at least one ATS, including iATS and cATS. **D-F.** Transcriptional probability broken down by proportion with ATS numbers from one to four (see legend top right). **G-I.** Transcriptional output measured as total number of nascent transcripts at cATS per cell row. **J-L.** Transcriptional output measured as total number of cellular mRNAs (not rachis) per cell row. Germ cell boundaries were determined from MATLAB-generated Voronoi cells centered in the nucleus (see Methods). **M-0.** Transcriptional output measured as number of mRNAs within each cell (not rachis) as a function of position.

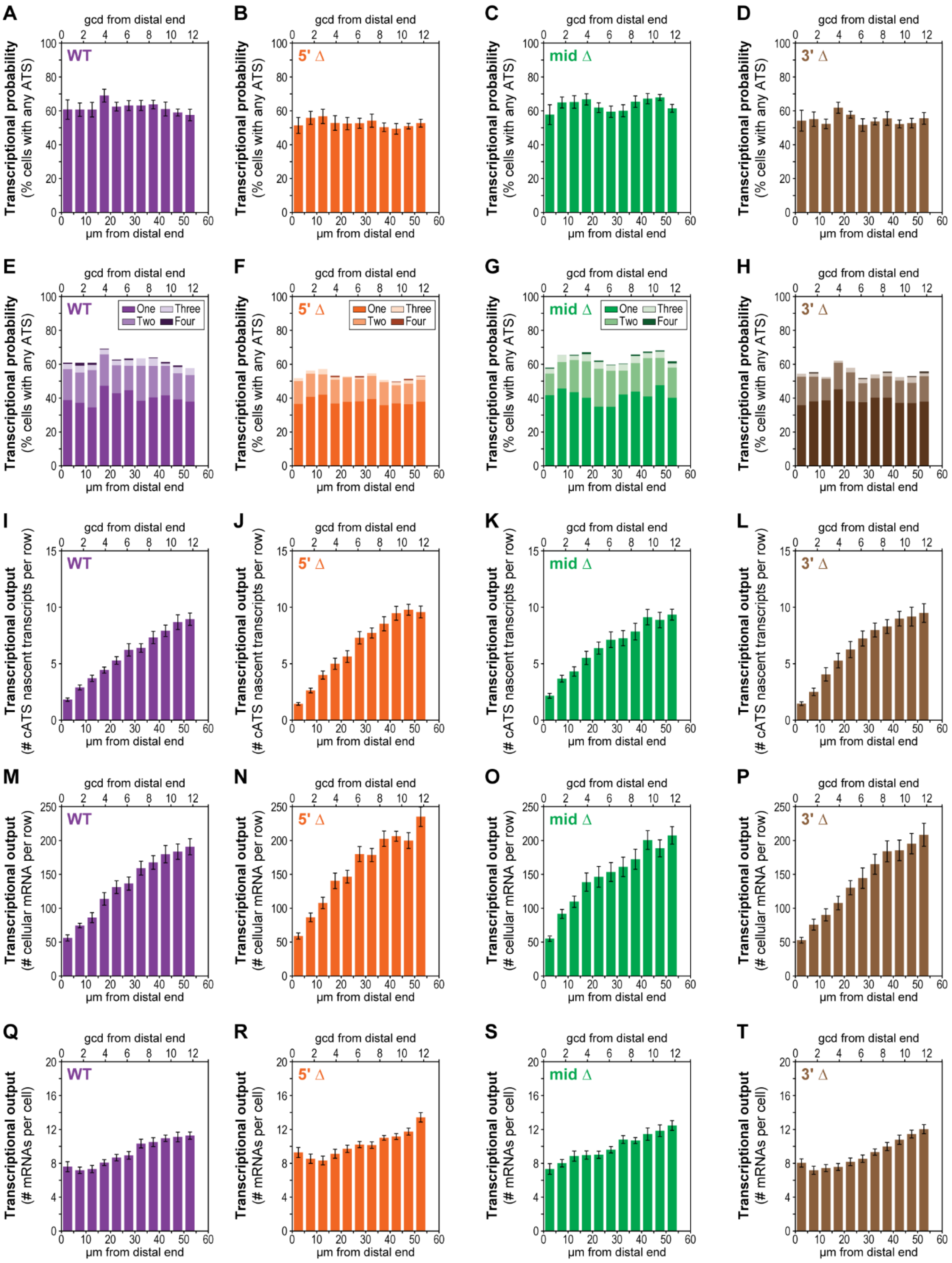


**Figure S9: Extended dataset for Figure 5. A-Q.** Transcriptional analysis of wildtype (WT) and three 1 kb *mpk-1b* intron deletions; see Figure 5A for sites of intron deletions. x-axis, gcd (top) and µm (bottom) from distal end.

Standard error bars shown. **A-D.** Transcriptional probability measured as percentage of cells with at least one ATS, including iATS and cATS. Number cells scored: wildtype, n = 2688; 5' Δ, n = 3361; mid Δ, n = 3434; and 3' Δ, n = 3302. **E-H.** Transcriptional probability broken down by proportion with ATS numbers from one to four (see legend top right). **I-L.** Transcriptional output measured as total number of nascent transcripts at cATS per cell row. **M-P.** Transcriptional output measured as total number of cellular mRNAs (not rachis) per cell row. Germ cell boundaries were determined from MATLAB-generated Voronoi cells centered in the nucleus (see Methods). **Q-T.** Transcriptional output measured as number of mRNAs within each cell (not rachis) as a function of position.

Supplemental Tables:

Table S3: Strains used in study

| **Strain Name** | **Genotype** | **Name in text** |
|---|---|---|
| N2 | wildtype | WT |
| JK6166 | *mpk-1 (q1069)/qC1 [qIs26]* III | *mpk-1b (fs)* |
| JK6033 | *mpk-1 (q1030)* III | 5' Δ |
| JK6130 | *mpk-1 (q1026)* III | mid Δ |
| JK6035 | *mpk-1 (q1040)* III | 3' Δ |
| JK6164 | *mpk-1 (q1084)* III | large Δ (*mpk-1* intron control) |
| JK6402 | *lag-3 (q1200)/qC1 [qIs26]* III | *lag-3* intron control |

Table S4: crRNAs used in study

| Name | Sequence (5' → 3') |
|---|---|
| mpk-1 intron 1.1 | ttttggaccgtctcgcgaaa |
| mpk-1 intron 1.2 | gctggaaaacctctacaaga |
| mpk-1 intron 1.3 | gcccattgtgctcccattat |
| mpk-1 intron 1.4 | gtattcctttgggaaccccg |
| mpk-1 intron 1.5 | gatttcctcgtcaacaagtc |
| mpk-1 intron 1.6 | gtgggaaactggaaatcacg |
| mpk-1b crRNA | AATGCTAAACCACCATCGAA |
| lag-3_1 | gacatttacactgaaaagtg |
| lag-3_2 | gtaggaacctgagagactcg |

Table S5: smFISH probe concentrations

| Probe Name | Number of probes | Fluor | Dilutions from stock | Final concentration |
|---|---|---|---|---|
| *mpk-1b* exon (original) | 48 | Far red 610 | 1:20 | 0.125 $\mu M$ |
| Intron *mpk-1b* (original) | 48 | Quasar 570 | 1:10 | 0.25 $\mu M$ |
| *mpk-1* intron 2 (swapped) | 48 | Far red 610 | 1:10 | 0.25 $\mu M$ |
| *mpk-1* exon 2 (swapped) | 48 | Quasar 570 | 1:20 | 0.125 $\mu M$ |
| *lag-3* intron | 48 | Far red 610 | 1:10 | 0.25 $\mu M$ |
| *lag-3* exon | 47 | Quasar 570 | 1:20 | 0.125 $\mu M$ |
| *lag-1* intron | 48 | Quasar 670 | 1:10 | 0.25 $\mu M$ |
| *lag-1* exon | 48 | TAMRA dye | 1:10 | 0.25 $\mu M$ |

Reference for supplemental legends:

Lee C, Sorensen EB, Lynch TR, Kimble J. 2016. C. elegans GLP-1/Notch activates transcription in a probability gradient across the germline stem cell pool. *Elife* **5**:e18370. doi:10.7554/elife.18370
